# Immunoglobulin superfamily receptor Junctional adhesion molecule 3 (Jam3) requirement for melanophore survival and patterning during formation of zebrafish stripes

**DOI:** 10.1101/2021.03.01.433381

**Authors:** Dae Seok Eom, Larissa B. Patterson, Raegan R. Bostic, David M. Parichy

## Abstract

Adhesive interactions are essential for tissue patterning and morphogenesis yet difficult to study owing to functional redundancies across genes and gene families. A useful system in which to dissect roles for cell adhesion and adhesion-dependent signaling is the pattern formed by pigment cells in skin of adult zebrafish, in which stripes represent the arrangement of neural crest derived melanophores, cells homologous to melanocytes. In a forward genetic screen for adult pattern defects, we isolated the *pissarro* (*psr*) mutant, having a variegated phenotype of spots, as well as defects in adult fin and lens. We show that *psr* corresponds to *junctional adhesion protein 3b* (*jam3b*) encoding a zebrafish orthologue of the two immunoglobulin-like domain receptor JAM3 (JAM-C), known for roles in adhesion and signaling in other developing tissues, and for promoting metastatic behavior of human and murine melanoma cells. We found that zebrafish *jam3b* is expressed post-embryonically in a variety of cells including melanophores, and that *jam3b* mutants have defects in melanophore survival. Jam3b supported aggregation of cells *in vitro* and was required autonomously by melanophores for an adherent phenotype *in vivo*. Genetic analyses further indicated both overlapping and non-overlapping functions with the related receptor, Immunoglobulin superfamily 11 (Igsf11) and Kit receptor tyrosine kinase. These findings suggest a model for Jam3b function in zebrafish melanophores and hint at the complexity of adhesive interactions underlying pattern formation.

## 1. Introduction

Adhesive interactions among cells and between cells and local extracellular matrices are essential to tissue development and homeostasis. Regulation of such interactions is likely to be especially important in highly migratory populations, like pigment cells derived from the neural crest, the progenitors of which must initiate migration at specific times, traverse long distances through heterogenous tissue environments, and then settle in precise arrangements relative to one another and other cell types (Kelsh et al., 2009).

Roles for cell–cell and cell–extracellular matrix interactions in pigment cell development are suggested by classical and more recent observations (Erickson and Perris, 1993; Haass et al., 2005; McKeown et al., 2013; Pinon and Wehrle-Haller, 2011). For example, pigment cells of some species develop persistent contacts with one another (Hamada et al., 2014; Parichy, 1996; Twitty, 1945) or express different cell–cell adhesion molecules as they mature (Nishimura et al., 1999). Likewise mutations or manipulations that impact cell–cell interactions can influence morphogenetic behaviors *in vitro* in ways likely to impact pattern formation in the animal (Garcia-Lopez et al., 2005; Inaba et al., 2012; Rao et al., 2016). Ontogenetic changes in extracellular matrices similarly correlate with migratory pathway choice, and alterations in pigment cell–matrix interactions affect motility *in vitro* (Dilshat et al., 2021; Parichy et al., 2006) as well as migration and colonization of skin *in vivo* (Haage et al., 2020; Ustun et al., 2019). Elucidating roles for adhesive interactions has been challenging, however, likely owing to redundant or compensatory functions within and between gene families (Williams et al., 2018).

Genetic and cellular approaches using zebrafish offer opportunities to further define roles for adhesion in pigment cell development and pattern formation. Skin pigment cells of zebrafish and other ectothermic vertebrates arise from neural crest cells, either directly or indirectly via latent progenitors, similar to melanocytes of birds and mammals (Adameyko et al., 2009; Budi et al., 2011; Dooley et al., 2013; Kelsh et al., 2009). Zebrafish pigment cells include black melanophores that contain melanin and are homologous to avian and mammalian melanocytes, and also yellow or orange xanthophores with pteridine or carotenoid pigments, and iridescent iridophores, the appearance of which depends on stacks of reflective guanine crystals (Gur et al., 2020; Parichy, 2021; Saunders et al., 2019; Schartl et al., 2016). Unlike birds and mammals, pigment cells of ectothermic vertebrates retain their pigments intracellularly rather than transferring them to keratinocytes; pigment patterns thus reveal the arrangements and differentiation states of the pigment cells that comprise them.

In adult zebrafish, pigment cells within the hypodermis of the skin are responsible for a striped pattern (Aman and Parichy, 2020; Hirata et al., 2003). Melanophores and sparsely arranged, bluish iridophores are arranged in dark stripes, whereas xanthophores and densely arranged, yellowish iridophores form light “interstripes” (**Fig. 1A,B**, *wild-type*) (Gur et al., 2020; Patterson and Parichy, 2019). Additional pigment cells are found more superficially on scales, where melanophores provide a dark cast to the dorsum.

**Fig. 1.**
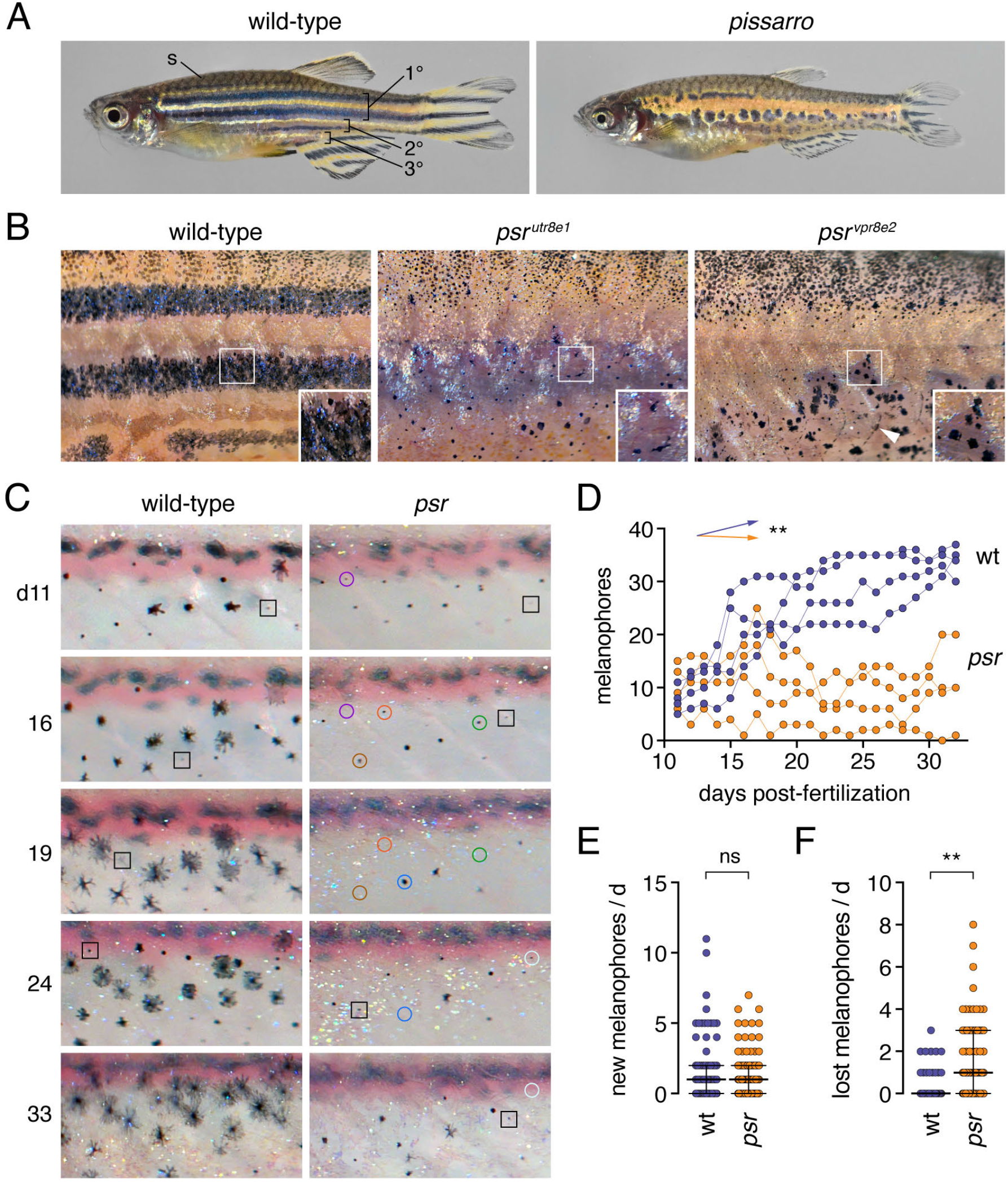
Phenotypes and pattern ontogeny of wild-type and *pissarro* (*psr*) mutant zebrafish. (**A**) Wild-type fish adult with primary, secondary and tertiary pattern elements and scale (s) melanophores labeled. *psr* mutants (here *psr*^*vp8re2*^) have disrupted melanophore stripes and fins that are short relative to body size. (**B**) Higher magnification images illustrating disorganized melanophores and iridophores in wild-type fish or homozygous mutants for different *psr* alleles. In *psr*^*vpr8e2*^ some melanophores were apparent on ventral scales (arrowhead), typical of the wik genetic background in which the allele was maintained. Insets illustrate well-spread appearance of most stripe melanophores in wild-type and heterogeneity of melanophore morphologies in *psr* mutant alleles. (**C**) Repeated daily imaging of representative wild-type and *psr*^*utr8e1*^ mutant larvae through adult pigment pattern formation. Days at left indicate days post-fertilization. Boxes indicate newly differentiating melanophores. Colored circles indicate melanophores lost from the pattern. These same fish are shown in Movies S1, S2. (**D**) Total numbers of melanophores increased in each of 4 repeatedly imaged wild-type larvae but remained at initial levels or decreased in each of 4 *psr* mutants in ventral portions of 3 segments. Differences in melanophore numbers were reflected in a significant genotype x day interaction after controlling for variation among individuals (**, *F*_1,6_=23.7, *P*=0.0028; arrows indicate slopes of partial regression coefficients). (**E**) Numbers of newly differentiating melanophores per day did not differ between wild-type and *psr* mutants after controlling for variation among individuals (ns, *F*_1,6_=0.3, *P*=0.6099). (**F**) Melanophores were, however, significantly more likely to be lost from *psr* mutants than wild-type after controlling for variation among individual fish (**, *F*_1,6.2_=16.5, *P*=0.0063; counts of melanophores were square root-transformed for analysis to correct for heterogeneity in residual variance between genotypes). Bars in E and F are medians ± interquartile range.

The zebrafish adult pattern forms during the larva-to-adult transformation as pigment cell precursors, derived from multipotent progenitors in the peripheral nervous system, migrate to the hypodermis (Budi et al., 2011; Dooley et al., 2013; Singh et al., 2016). The first cells to arrive differentiate as iridophores of a primary interstripe, whereas later arriving cells differentiate as yellowish interstripe iridophores, or as melanophores that settle further dorsally and ventrally in primary stripes (**Fig. 1A,B**) (Parichy et al., 2009; Patterson and Parichy, 2013). Subsequently, unpigmented xanthophores—residing already in the hypodermis—begin to mature in the interstripe (McMenamin et al., 2014). Finally, additional progenitor-derived cells differentiate as sparesely arranged bluish iridophores in the stripes (Gur et al., 2020). Fish grow substantially during pattern formation, and as they do, these events are reiterated to form secondary and tertiary interstripes and stripes; more superficially, additional pigment cells populate scales, which also develop during this period. The first indications of adult pattern formation occur at ~1.5 weeks post-fertilization (~5 mm standardized standard length, SSL), primary and secondary stripes and interstripes and scale pigment cells form by ~4 weeks (13 SSL), and adult pattern is completed by ~8 weeks (~20 SSL) though absolute timing depends on rearing conditions (McMenamin et al., 2016; Parichy et al., 2009).

Forward genetic screens for zebrafish mutants have uncovered genes whose products function in, or depend on, cell adhesion during adult pigment pattern formation. For example, several genes, including *leopard*, encode gap junction or tight junction proteins that are expressed and required by pigment cells, and when activities of these genes are perturbed, spots form instead of stripes (Fadeev et al., 2015; Irion et al., 2014; Mahalwar et al., 2016; Usui et al., 2019; Watanabe et al., 2006; Watanabe et al., 2016). A different class of adhesion molecule is represented by Coxsackie- and Adenovirus Receptor (CAR) and Junctional Adhesion Molecule (JAM) family genes, which encode dimeric cell surface adhesion receptors having extracellular regions with two immunoglobulin (Ig)-like domains and intracellular regions that interact with PDZ-containing proteins that function in scaffolding and signaling (Ebnet, 2017; Ebnet et al., 2004; Kummer and Ebnet, 2018; Rathjen, 2020; Schreiber et al., 2014). CAR and JAM family members participate in homophilic and heterophilic interactions in *cis* and *trans* to regulate cell-cell contacts, junctional adhesion, migration, motility, proliferation and polarity across a wide range of cell types. In zebrafish, the CAR family member Immunoglobulin superfamily 11 (Igsf11) allows for homophilic interactions, and promotes migration and survival of melanophores. *seurat* mutant zebrafish have missense substitutions in Igsf11 and develop spots similar to mutants for gap junction proteins (Eom et al., 2012).

Here, we identify a new cell adhesion molecule with essential roles in pigment cell behavior and patterning. We show that a different spotted mutant of zebrafish, *pissarro*, results from lesions in *junctional adhesion molecule 3b* (*jam3b*), encoding a JAM family receptor. Jam3b allowed for adhesive of heterologous cells and was essential in melanophores for an adherent morphology and survival. We further identified genetic interactions that suggest overlapping and non-overlapping roles of *jam3b* with *igsf11* and a gene encoding Kit receptor tyrosine kinase, *kita*. These findings reveal a new gene family with roles in the development of skin pigment pattern, and point to roles for adhesive interactions during morphogenesis and maturation of pigment cells *in vivo*.

## 2. Materials and Methods

### 2.1. Fish care, lines and crossing

Fish were reared under standard conditions (14L:10D at ~28°C) and staging followed (Parichy et al., 2009). Fish were fed marine rotifers, brine shrimp and flake food. Rearing and experimental analyses followed protocols approved by Institutional Animal Care and Use Committees of University of Texas, University of Washington, and University of Virginia (current protocol #4286). Fish were euthanized by overdose of MS-222.

Fish lines used or generated were: *psr/jam3b*^*utr8e1*^, *psr/jam3b*^*vp8re2*^, *psr/jam3b*^*vp8re3*^, *igsf11*^*vp15rc4*^, *igsf11*^*vp15rc5*^, TgBAC(*jam3b:jam3b-mCherry*)^*vp36rTg*^ (this study); *gja5b*^*t1*^ (*leopard*) (Watanabe et al., 2006); *igsf11*^*wp15e1*^ (Eom et al., 2012); *kita*^*b5*^ (*sparse*) (Parichy et al., 1999); *ltk*^*j9s1*^ (*primrose*) (Lopes et al., 2008); TgBAC (*tyrp1b:PALM-mCherry*)^*wprt11Tg*^, TgBAC (*aox5:PALM-GFP*)^*wprt12*^ (McMenamin et al., 2014); Tg (*pnp4a:palm-EGFP*)^*wprt22Tg*^; Tg(*kdrl:GFP*) (Choi et al., 2007); Tg(*mitfa:EGFP*)^*w47*^ (Budi et al., 2011; Curran et al., 2010); Tg(*actb2:EGFP*) (provided by K. Poss).

Crosses were performed by natural spawning or *in vitro* fertilization and genotypes verified by Sanger sequencing for allele-specific lesions (*psr*^*utr8e1*^: 5’- TTTTAATTCCTGATGTGTCAACAAG-3’, reverse 5’-ATTTCTGGTCATTGGGTGCT3’; *psr*^*vp8re2*^: 230 bp amplicon, 5’-CTTCGGAATGACGTTAAGATGA-3’, CCCATCTAGGGTGTAAGTGGAG-3’).

### 2.2. *Isolation and molecular cloning of* psr *mutant alleles*

Fish were mutagenized with N-ethyl-N-nitrosourea using standard methods (Solnica-Krezel et al., 1994) and F2s screened for novel phenotypes following early pressure gynogenesis (Johnson et al., 1995). *psr*^*utr8e1*^ was isolated in the AB^wp^ background, derived from University or Oregon AB* and maintained as in inbred stock since 1998. *psr*^*utr8e1*^ subsequently mapped by crossing to fish of the inbred wik strain and backcrossing resulting heterozygous F1s to homozygous *psr*^*utr8e1*^. Informative markers were identified by Sanger sequencing of AB^wp^ and wik backgrounds and candidate genes within a critical recombination interval were sequenced by Sanger sequencing. Additional sequencing of *psr*^*vp8re2*^ was obtained by low-coverage whole genome resequencing of a single mutant individual, with subsequent validation by Sanger sequencing.

### 2.3. Reverse transcription polymerase chain reaction (RT-PCR) analysis of gene expression in pigment cells

For assaying gene expression by RT-PCR adult skins were first dissociated by incubating for 10 min in calcium and magnesium free phosphate buffered saline (PBS) containing trypsin, followed by gentle pipetting. Individual cells were then picked manually under a stereomicroscope, identifying cells by pigment phenotype (melanophores: melanin; xanthophores: yellow-orange color and autofluorescence under epiflourescence with 488 nm green filter set; iridophores, iridescence under brightfield incident illumination and reflectance under epiflourescent illumination with both 488 nm / green and 546 nm / red filter sets). For each cell type ~100 cells were collected; portions of whole fins were additionally collected. Total RNAs were isolated by RNeasy Protect Mini Kit (Qiagen) and cDNAs synthesized with oligo-dT priming using SuperScript III Cells Direct cDNA Synthesis System (Thermo). Primers pairs were designed to span introns for assessing genomic contamination, targeting *jam3b* (5’- AAGGAGGAAATCCCGCTAGA-3’, 5’-AAGAGGACCATCACCACCAC-3’), *igsf11* (5’- TCTGATCGCGGGCACCATCG-3’, 5’-TAGGTGTTGGTGGACGTCAGAGTG-3’), and loci preferentially expressed across pigment cell classes (Eom et al., 2012; Saunders et al., 2019). Amplifications were performed using Taq polymerase with 40 cycles of 95°C for 30 s, 60°C for 30 s, 72°C for 30 s.

### 2.4. Recombineering, transgenesis and re-analysis of single cell RNA-sequencing data

To generate a fluorescent reporter of *jam3b* expression and localization, bacterial artificial chromosome (BAC) CH73-346N23 containing *jam3b* and upstream regulatory sequence was identified through the Ensembl genome browser (release GRCz10) and recombineered within the vector backbone to contain inverted *Tol2* repeats followed by recombineering at the C-terminus to replace the native terminator codon with an mCherry cassette containing an SV40 polyadenylation sequence (Sharan et al., 2009; Suster et al., 2011). For transgenesis, the recombineered BAC, *jam3b*:Jam3b-mCherry, and *Tol2* mRNA was injected into zebrafish embryos at the 1-cell stage (Suster et al., 2011; Suster et al., 2009) and stable carriers isolated by standard methods.

An additional construct for expressing Jam3b *in vivo* was generated using Tol2Kit plasmids for Gateway construction (Kwan et al., 2007) with a p5E element consisting of 2.2 kb upstream of the *melanocyte inducing transcription factor a* (*mifta*) transcription start site to drive expression in melanophores (Eom et al., 2012), a pME element consisting of Kozak sequence with wild-type *jam3b* cDNA linked by peptide breaking 2a sequence to nuclear localizing venus, and a p3E SV40 polyA signal. The resulting construct, mitfa:jam3b-2a-nVenus was co-injected with *Tol2* mRNA and analyzed in F0 mosaic individuals.

Assessment of *jam3b* expression in a previously published single cell RNA-sequencing dataset (GEO Accession: GSE131136) used Monocle 3 (Cao et al., 2019; Saunders et al., 2019).

### 2.5. Immunohistochemistry

Immunostaining for zebrafish Igsf11 used a polyclonal antiserum generated in mouse and characterized previously (Eom et al., 2012). Co-labeling with *jam3b*:Jam3b-mCherry was performed on melanophores isolated as described above from 7.5 standardized standard length (SSL) larvae (Parichy et al., 2009). Melanophores were plated onto poly-D-lysine coated glass-bottom dishes and lightly fixed in 4% paraformaldehyde in PBS. After blocking anti-Igsf11 antiserum was used at a concentration of 1:500 and detected with goat anti-mouse Alexa 488 secondary antibody (ThermoFisher).

### 2.6. S2 cell aggregation assays

*Drosophila* S2 cells were maintained in Schneider’s medium (ThermoFisher) and transfected with constructs for fusion protein cDNAs cloned into the pJFRC (Janelia Reporter Construct) backbone using Actin-EGFP as a control. Constructs generated with standard methods were pJFRC-Jam3b-mCherry, pJFRC-Jam3b(A70E)-mCherry, and pJFRC-Igsf11-EGFP to label Jam3b or Igsf11 protein with mCherry or EGFP, respectively. Cells were transfected with Lipofectamine (ThermoFisher) using the manufacturer’s recommended protocol. Transient Actin, Jam3b, Jam3b(A70E), or Igsf11 expression was confirmed by visualizing tagged fluorophores by widefield epiflourescence microscopy. Following transfection, cells were collected by centrifugation, resuspended by gentle pipetting, and aliquoted into 24-well culture dishes at 6 x 10^4^ cells/ml. Dishes were rotated on a gyratory rotator at 80 rpm for 2 h and randomly selected areas imaged every 30 min from each culture. Aggregation was assessed by calculating numbers of transfected cells found in clusters relative to total number of transfected cells present.

### 2.7. Imaging

Images of whole fish or gross pattern features were captured on a Nikon D-810 digital single lens reflex camera with MicroNikkor 105 mm macro lens; an Olympus SZX-20 stereomicroscope with Zeiss Axiocam HR camera; or on a Zeiss AxioZoom stereomicroscope or Zeiss Axio Observer inverted microscope, each equipped with wide-field epifluorescent illumination and Zeiss Axiocam 506 monochrome and color cameras with ZEN blue software. High resolution fluorescence imaging used a Zeiss LSM800 and Zeiss LSM880 Observer inverted laser confocal microscopes equipped with GaASP detectors and Airyscan.

To assess gross features of pigment cell behaviors during adult pattern development, individually reared *psr* mutant and wild-type siblings were imaged daily between 6.5–8.0 SSL using an Olympus SZX-20 stereomicroscope with Zeiss Axiocam HR camera (Eom et al., 2012; Parichy and Turner, 2003b).

Time-lapse imaging of *mitfa*:GFP+ melanoblasts was performed with wild-type and *psr* mutant trunks explanted to glass bottomed dishes and followed for 15 h at 30 min intervals on a Zeiss AxioObserver inverted microscope with wide-field epifluorescent illumination and Zeiss Axiocam 506 monochrome camera (Budi et al., 2011; Eom et al., 2012).

For repeated imaging of individual larvae, images captured on successive days were scaled isometrically relative to one another and aligned by eye in Adobe Photoshop 2021. Regions of interest for assessing hypodermal (stripe) melanophore development were defined for three segments immediately posterior to the vent extending ventrally from horizontal myoseptum to ventral margin of the flank, thereby excluding scale melanophores, which occur further dorsally. Within this region, melanophores were censused and newly differentiating melanophores or unambiguously lost melanophores recorded; melanophores leaving or entering the region of interest were not included in counts of new or lost cells.

For comparing melanophores complements across *psr* mutant alleles of adult fish, ventral regions of interest were defined across five segments, extending anteriorly from the posterior margin of the anal fin so as to include the region where the primary ventral melanophore stripe would form normally (ventrally from the horizontal myoseptum to a point halfway between myseptum and ventral margin of the flank). Fish of the wik genetic background sometimes exhibit non-hypodermal scale melanophores in ventral regions and such cells were excluded from analyses.

For display, images were adjusted for levels and color balance using Zeiss ZEN software or Adobe Photoshop 2021. The ZEN Extended Focus module was used for projections across focal planes for some brightfield or fluorescence images. Corresponding images across genotypes or treatments were adjusted concordantly.

### 2.8. CRISPR/Cas9 mutagenesis

To induce germline or somatic (mosaic) mutations we used CRISPR/Cas9 mutagenesis by injecting 1-cell stage embryos with 200 ng/*μ*l T7 sgRNA and 500 ng/*μ*l Cas9 protein (PNA Bio) as described (Shah et al., 2015). New alleles of *igsf11* were produced by targeting a site within exon3 (igsf11ex3-C1: 5’-GGCACTTGTGCTGGGCATGG-3’; screening primers: 5’- TTCAGTTTAGCCAATCACATGAA-3’, 5’-TCAGGCCGATCACTCCTATATT-3’) and were then bred to homozygosity. For somatic mutagenesis of *jam3b* we targeted a site in exon 2 (5’- GgCGACGAATTCTAAACCGT-3’), which yielded phenotypes indistinguishable from previous isolated germline alleles of *jam3b*.

### 2.9. Statistical analyses

Analyses of quantitative data were performed in JMP14 statistical analysis software (SAS Institute) with parametric model assumptions verified by inspection of residuals and transformations of original data as warranted to normalize variances across groups. In analyses of repeated daily imaging of individual larvae, mixed model analyses of variance were used to control for differences among individuals (random effect) within genotypes (treated as a fixed effect) to avoid pseudoreplication.

## 3. Results

### 3.1. *Defective adult stripe formation in* pissarro *mutants*

In a forward genetic screen for ENU-induced mutations affecting post-embryonic development, we isolated the recessive and homozygous viable *pissarro* mutant (**Fig. 1A**), named for the Danish-French impressionist and pointillist Camille Pissarro, who worked alongside Georges Seurat. Initial screening was performed in the inbred AB^wp^ genetic background, yielding a single allele, *psr*^*utr8e1*^, that was maintained in this same genetic background. Two additional ENU-induced alleles, *psr*^*vp8re2*^ and *psr*^*vp8re3*^, were subsequently isolated by non-complementation screening against the original allele, using ENU-mutagenized fish of the wik genetic background, in which these new alleles were subsequently maintained.

Adult fish homozygous for *psr* mutant alleles had disrupted patterns with hypodermal melanophores that were heterogeneous in shape and size (**Fig. 1B**). *psr*^*ut8re1*^ and *psr*^*vp8re2*^ homozygotes had ~17% and ~48% as many hypodermal melanophores as wild-type fish (**Fig. S1A** and see below). Iridophore morphologies were similar to wild-type, with yellowish and densely packed iridophores in most regions lacking melanophores, and bluish, sparsely arranged iridophores where melanophores were present (**Fig. S1B**). Dense iridophores, typical of interstripe regions, were spread more widely over the flank than in the wild type, similar to other mutants with reduced numbers of melanophores (Frohnhofer et al., 2013; Gur et al., 2020). Yellow-orange xanthophores had grossly normal morphologies and were associated with dense iridophores, as in wild-type and other melanophore-deficient mutants. Scale melanophore complements were not patently different from wild-type (**Fig. 1A,B**). Fin pigment patterns were somewhat irregular, though fins were themselves smaller and fin bones (lepidotrichia) shorter than wild-type (**Fig. 1A**, **Fig. S2A,B**). *psr* mutant adults also developed lens opacities (**Fig. S2C,D**).

To better understand the ontogeny of adult pigment pattern in *psr*, we followed pigment cells in larvae imaged daily through juvenile stages (**Fig. 1C**; **Movies S1, S2**). In wild-type, melanophore numbers in a ventral regions of three representative segments increased through pattern formation whereas in *psr* mutants melanophore populations failed to increase and in some cases decreased (**Fig. 1D**), the latter yielding phenotypes more severe than observed typically. Such increased severity likely reflects an enhancement to the normal sensitivity of pattern formation to variation in somatic growth rate or rearing conditions (Parichy et al., 2009), as observed for some other mutants as well (Eom et al., 2012; Parichy and Turner, 2003b).

By following the differentiation and morphogenesis of individual cells, we found that new melanophores appeared in *psr* mutants at rates and at locations similar to wild-type (**Fig. 1C,E**). Displacements of immature melanophores were similar to wild-type, as were movements of melanoblasts observed by short-duration time-lapse imaging (**Movies S3, S4**). By contrast, melanophores were significantly less likely to persist in *psr* mutants than in wild-type, with medians of 1 and 0 melanophores lost per day, respectively, from regions of interest (means±SD = 1.4±1.8, 0.3±0.6) (**Fig. 1F**), indicative of cell death and extrusion (Eom et al., 2012; Lang et al., 2009; Quigley et al., 2005). Dense iridophores came to occupy regions newly devoid of melanophores, as seen in other contexts (Gur et al., 2020), thus generating an irregular pattern overall (**Movies S1, S2**). During later development, adult *psr* mutants also exhibited melanized debris not observed in wild-type fish, consistent with an increased likelihood of melanophore death even after pattern formation would normally be completed (**Fig. S1C**). These observations show that *psr* normally promotes the survival of melanophores during and after stripe development. Variegated patterns of *psr* mutants likely result from this inappropriate melanophore death and cascading effects of altered melanophore numbers and positions on other pigment cells that normally interact with melanophores (Gur et al., 2020; Patterson and Parichy, 2019; Watanabe and Kondo, 2015).

### 3.2. psr *encodes Junctional adhesion molecule 3b*

To identify the *psr* gene we crossed *psr*^*ut8re1*^ homozygotes to fish of the inbred wik genetic background and then mapped the phenotype on a panel of 3040 backcross individuals, which defined a critical interval of ~300 kb on Chr21 containing 15 genes (**Fig. 2A**). Sanger sequencing of 8 candidates for *psr*^*utr8e1*^ revealed a missense substitution (A96E) not present in the unmutagenized backround in the fourth exon of *junctional adhesion molecule* 3b (*jam3b*) (**Fig. 2B**). Low-coverage whole-genome re-sequencing of *psr*^*vp8re2*^, verified by Sanger sequencing, likewise revealed a premature stop codon (Q174X) in the region encoding the second Ig-domain of Jam3b, but did not identify mutations in other candidate genes. Finally, targeted Sanger sequencing identified a novel missense substitution (C234W) in a third phenotypically similar allele, *psr*^*vp8re3*^, used for gene identification but not developmental analyses. These findings demonstrate the molecular identity of *psr*, referred to hereafter as *jam3b*, and in so doing identify another Ig-superfamily member with roles in pigment pattern formation, potentially having roles similar to *igsf11* (Eom et al., 2012).

**Fig. 2.**
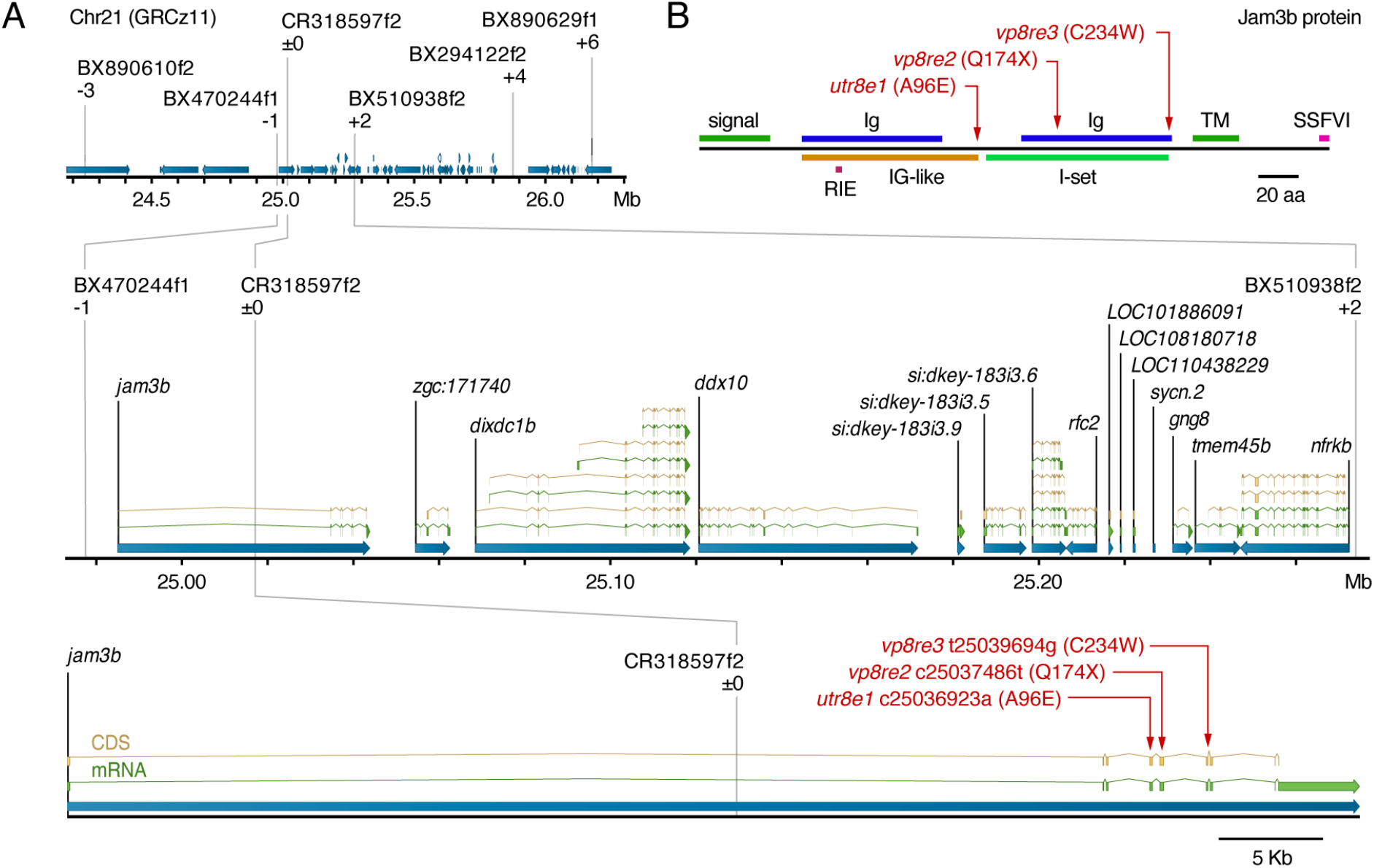
Positional cloning of *psr*. (**A**) Region of Chr21 to which *psr* was mapped using polymorphisms identified in AB^wp^ and wik backgrounds. Informative markers and recombinants relative to *psr*^*utr8e1*^ phenotype are shown above chromosome. Positions of causal substitutions below, relative to zebrafish whole genome assembly GRCz11 (nucleotides) or Jam3b protein (amino acids). (**B**) Amino acid substitutions in *psr* mutant alleles relative to Jam3b protein domains. RIE, homophilic interaction domain tripeptide; TM, transmembrane domain; SSFVI, type II PDZ-domain-binding motif (Ebnet, 2017; Ebnet et al., 2004; Kostrewa et al., 2001; Powell and Wright, 2012; Santoso et al., 2005).

Because *jam3b*^*utr8e1*^ and *jam3b*^*vp8re2*^ were isolated and maintained in different genetic backgrounds, we asked whether differences in severity of their phenotypes were specific to the lesions themselves or the backgrounds in which they were maintained. We therefore generated *jam3b*^*utr8e2*^/*jam3b*^*wp8re1*^ fish that we intercrossed for assessing each of the three possible mutant genotypes. These analyses did not reveal differences in melanophore complements or lengths of fin bones across allelic combinations, though lens opacities differed between alleles even in a common genetic background (**Fig. S1A, Fig. S2B,C**). These outcomes suggest that A96E (*jam3b*^*utr8e1*^) and Q174X (*jam3b*^*vp8re2*^) are functionally similar with respect to pigmentation and fin bone development, though other cell types may be more sensitive to differences between missense and presumptive null alleles.

### 3.3. jam3b *is expressed in post-embryonic pigment cell lineages and other cell types*

In zebrafish embryos *jam3b* is expressed by somitic and lateral plate mesoderm and functions in myocyte fusion, vasculogenesis, and hematopoiesis (Kobayashi et al., 2020; Powell and Wright, 2011, 2012). *jam3b* expression and function during post-embryonic stages has not been described. To assess expression and interpret possible function within melanophores, xanthophores, or iridophores, we isolated pigment cells of each class from adult fish and tested for the presence of transcripts by RT-PCR, revealing expression in each of the three pigment cell classes (**Fig. 3A**). For earlier stages of pigment pigment pattern formation and pigment cell lineages we inspected a previously published, single cell RNA-sequencing dataset (Saunders et al., 2019) which revealed *jam3b* expression in melanophores, xanthophores, iridophores, and pigment cell progenitors (**Fig. 3B**). Jam3b can bind homophilically in *trans* to Jam3b, but also interacts *in vitro* and during myocyte fusion *in vivo* with Jam2a, to which binding is stronger (Powell and Wright, 2011, 2012). scRNA-Seq revealed *jam2a* expression in melanophores, iridophores, and their progenitors, but in very few xanthophores (**Fig. 3B**).

**Fig. 3.**
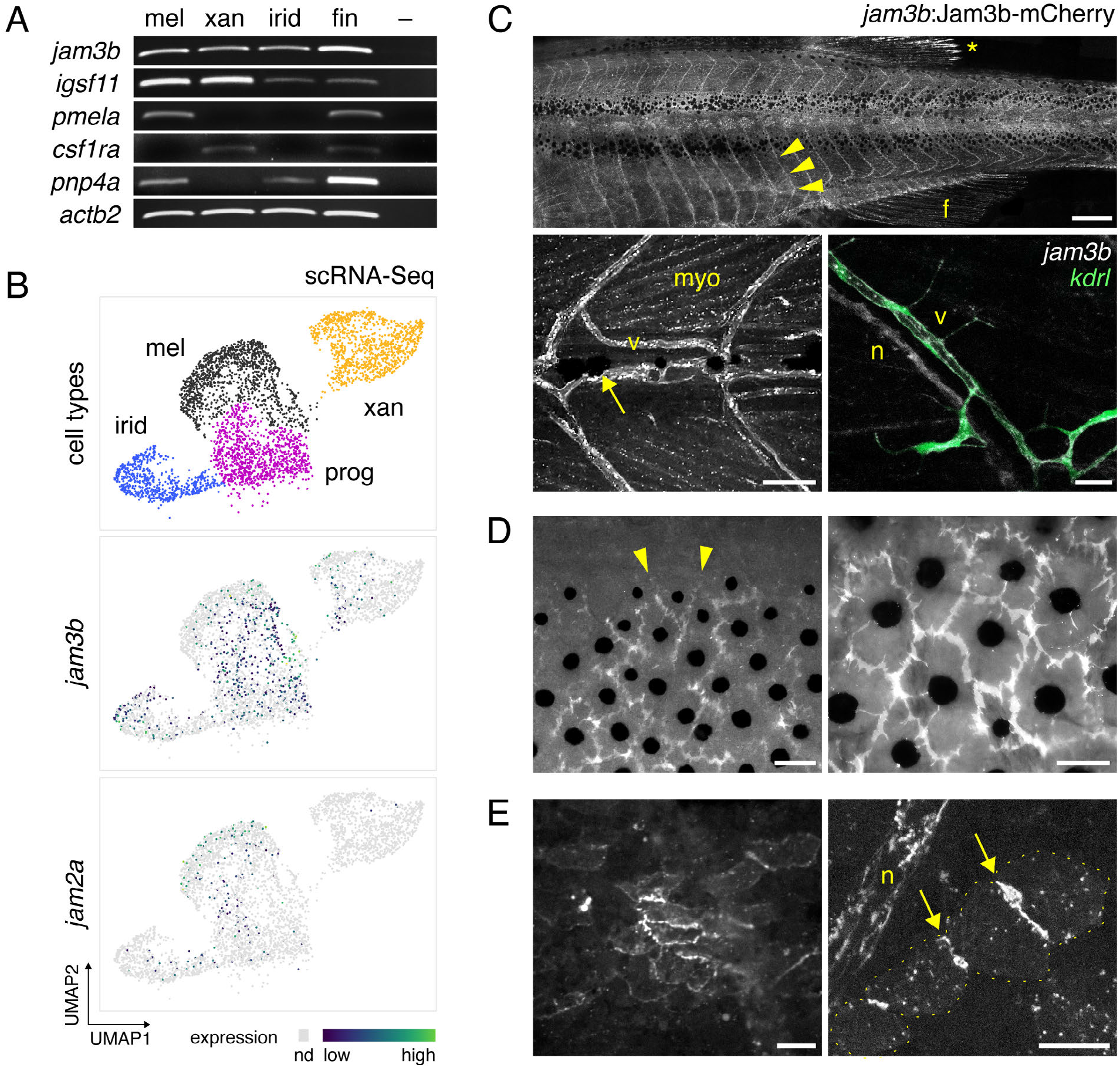
*jam3b* expression in pigment cells and other cell types of post-embryonic zebrafish. (**A**) RT-PCR for *jam3b* and related 2-Ig domain receptor gene *igsf11* in isolated melanophores (mel), xanthophores (xan), iridophores (irid), and whole fin tissue. –, no template control. *pmela* marks melanophores, *colony stimulating factor 1 receptor a* (*csf1ra*) marks xanthophores, and *purine nucleoside phosphorylase 4a* (*pnp4a*) most strongly marks iridophores and melanophores (Lang et al., 2009; Patterson and Parichy, 2013; Saunders et al., 2019). (**B**) *jam3b* and *jam2a* expression in melanophores, xanthophores, iridophores and post-embryonic pigment progenitor cells by single cell RNA-sequencing across states of maturation during adult pigment pattern formation. Cells in which transcripts were not detected (nd) are shown in light grey; the incidence of such cells is typical of loci expressed at moderate levels. *igsf11* transcripts were expressed at levels too low to detect by scRNA-Seq. (**C–E**) Expression of *jam3b*:Jam3b-mCherry. (**C**) *Upper,* Segmental expression (arrowheads) associated with vasculature (arrowheads) in a late larva (10 SSL). Diffuse expression is also evident in fin (f). *, highly reflective melanoleucophores (Lewis et al., 2019) of the dorsal fin, not representative of Jam3b-mCherry expression. *Lower left*, Expression in an earlier larva (6 SSL), with labeling of vasculature (v) as well as myocytes (myo). Arrow, early larval melanophore at horizontal myoseptum. *Lower right*, Jam3b-mCherry co-expression with endothelial marker *kdrl*:GFP (lower right). Additional Jam3b-mCherry was evident in presumptive Schwann cells of an adjacent nerve fiber (n) (Ebnet, 2017; Scheiermann et al., 2007). (**D**) Within mature stripes, Jam3b-mCherry accumulated at sites of apparent overlap between melanophores but typically did not extend to where melanophore membranes bordered the interstripe (arrowheads). Different regions are shown of the same fish (14 SSL) after partial contraction of melanin granules with epinephrine, using widefield epifluorescence to capture tissue context. (**E**) Low and high magnification images of interstripe iridophores (7 and 6 SSL, respectively), in which Jam3b-mCherry accumulated at sites of juxtaposition between cells (arrows). Yellow dots outline individual iridophores for clarity at right. Scale bars, 500 *μ*M, 50 *μ*M, 20 *μ*m (C), 50 *μ*M (D), 20 *μ*m (E).

To assess the tissue context of Jam3b expression, we recombineered a ~120 kb BAC, containing the *jam3b* gene body and ~18 kb upstream of the transcriptional start site, to express Jam3b fused at its carboxy terminus to mCherry, which we used to generate a stable transgenic line, TgBAC*(jam3b:jam3b-mCherry)*. In mutants *jam3b:jam3b-mCherry* failed to rescue phenotypes when crossed into *jam3b*^*utr8e1*^ or *jam3b*^*vp8re2*^ backgrounds. This outcome differed from another Jam3b transgene (below) and might reflect steric interference of C-terminally fused mCherry with essential interactions between cytosolic scaffolding proteins and the PDZ-protein binding motif of Jam3b, also located at the C-terminus (**Fig. 2B**) (Ebnet, 2017). The extent to which this or another factor might impact sub-cellular localization of Jam3b-mCherry remains uncertain. Nevertheless, expression of this transgene was detectable in an array of cell types, with particular abundance in vasculature and muscle cells of the myotomes, but also lens and basal cells of epidermis (**Fig. 3C**; **Fig. S3**), situated distant and across the skin basement membrane from melanophores in the hypodermis (Aman and Parichy, 2020; Hirata et al., 2003).

In pigment cells, Jam3b-mCherry was evident at edges of tiled melanophores within stripes (**Fig. 3D**), and where iridophores of interstripes were juxtaposed with one another (**Fig. 3E; Fig. S3**). Despite the presence of *jam3b* transcript in pigment cell precursors and xanthophores (**Fig. 3A,B**), we were unable to detect Jam3b-mCherry in migratory *mitfa*:GFP+ melanoblasts or in xanthophores. These disparities could result from insufficient time for the fusion protein to accumulate to detectable levels in rapidly developing melanoblasts, cell-type specific variation in post-transcriptional regulation in either cell type, as observed for other JAMs (Ebnet, 2017), or expression heterogeneity specific to the fusion protein itself.

### 3.4. *Melanophore lineage requirement for* jam3b *function*

To determine in which cell types *jam3b* is required during pigment pattern formation, we transplanted fluorescently labeled cells at the blastula stage between wild-type and mutant embryos and reared chimeras until juvenile pigment patterns developed. Where wild-type donor cells contributed to myotomes, *jam3b* mutant melanophores developed patterns typical of *jam3b* mutants (**Fig. 4A**). By contrast, where donor wild-type cells developed as melanophores or iridophores, their morphologies and pattern resembled those of the wild type (**Fig. 4B**). These results suggest that *jam3b* has an essential role within pigment cells.

**Fig. 4.**
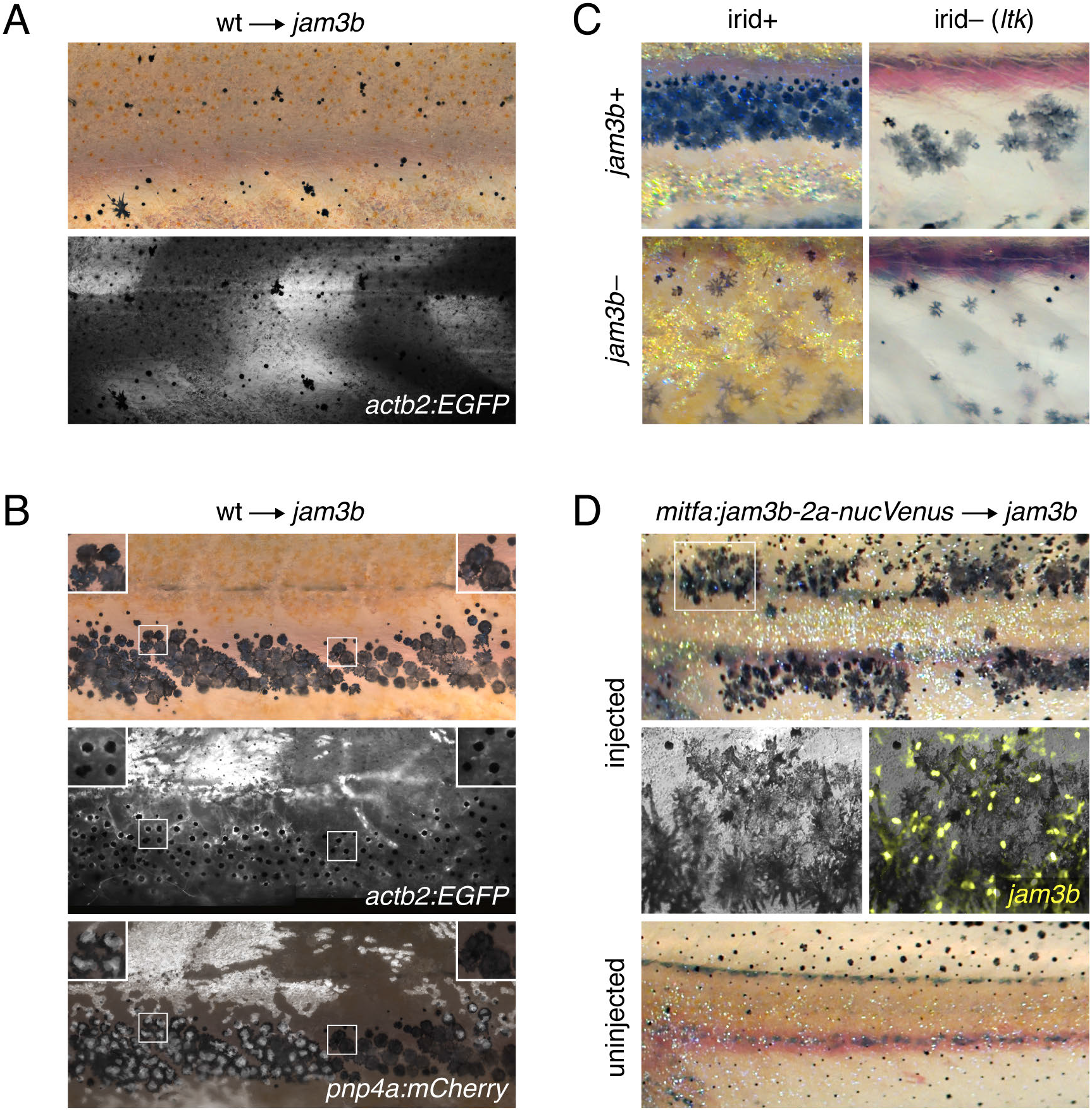
Cell-autonomy of melanophore requirement for Jam3b. (**A**) Fluorescently labeled wild-type cells that developed as myotome did not rescue the pigment pattern of *jam3b* mutant hosts. (**B**) Wild-type cells that developed as melanophores in *psr* mutant hosts exhibited wild-type morphologies and arrangements, whether or not these cells developed in the vicinity of wild-type iridophores (left and right details, respectively). Transplanted cells in these experiments were transgenic for *actb2:EGFP*, which is broadly expressed across a variety of cell types, but variegated in iridophores, and *pnp4a:PALM-mCherry*, which is expressed reliably in iridophores (Patterson and Parichy, 2013; Spiewak et al., 2018). Images are stitched from multiple tiles and melanosomes were contracted partially towards cell centers by epinephrine treatment to facilitate detection of *actb2:EGFP*. Informative chimeras examined, *n*=11. (**C**) In *ltk* mutant fish, iridophores are absent and fewer melanophores develop, resulting in spots rather than stripes (*upper right*) (Frohnhofer et al., 2013; Lopes et al., 2008; Patterson and Parichy, 2013). *jam3b* mutant melanophore phenotypes were similar whether fish were wild-type for *ltk* (irid+, *lower left*) or mutant for *ltk* (irid–, *lower right*). (**D**) *jam3b* mutants expressing wild-type Jam3b mosaically in melanophores had partially rescued melanophore morphologies and patterns (*upper panels*; uninjected control, *lower panel*). Note that *mitfa* and the *mitfa* regulatory region used here are expressed in melanophores and melanoblasts, as well as xanthophores; the binucleate state revealed by nucVenus expression is typical of mature, well-spread melanophores (Eom et al., 2012; Saunders et al., 2019).

Reciprocal interactions among all three classes of pigment cells during pattern formation (Frohnhofer et al., 2013; Gur et al., 2020; Patterson and Parichy, 2013, 2019; Watanabe and Kondo, 2015) raised the possibility that some effects of *jam3b* on melanophores might be indirect. Wild-type xanthophores did not develop in these chimeras, excluding the possibility that *jam3b* acts only through these cells to affect melanophore development. Because melanophores and iridophores arise from a common progenitor and developed together in chimeric fish, however, we considered that *jam3b* might function only in iridophores, with these cells influencing melanophores secondarily. Arguing against this possibility, wild-type melanophores in *jam3b* mutant hosts retained a wild-type morphology even when these cells were located at a distance from wild-type iridophores (**Fig. 4B**). Also consistent with melanophores requiring *jam3b* independently of iridophores, morphologies and arrangements of melanophores were similar between *jam3b* mutants and *jam3b* mutants constructed to lack iridophores owing to a mutation in *leukocyte tyrosine kinase* (*ltk*), which is normally required autonomously by iridophores for their development (**Fig. 4C**).

To further test a role for *jam3b* specific to melanophores we asked whether resupplying Jam3b to mutant melanophores can rescue their appearance and pattern. We constructed a transgene to drive wild-type Jam3b linked by peptide breading 2a sequence (Provost et al., 2007) to nuclear-localizing Venus, using regulatory elements of *microphthalmia inducing transcription factor a* (*mitfa*) (Eom et al., 2012). Injection of *psr* mutant embryos with this transgene yielded genetically mosaic animals with patches of *mitfa*:Jam3b-2a-nucVenus+ melanophores that had morphologies and arrangements resembling wild-type (**Fig. 4D**). A corresponding test for iridophores, using regulatory elements of *pnp4a* to express Jam3b in that lineage (Gur et al., 2020; Patterson and Parichy, 2013), failed to rescue the appearance or pattern of melanophores. Together these findings suggest that *jam3b* acts within melanophores to promote adult pigment pattern development.

### 3.5. *Jam3b promotes aggregation of S2 cells* in vitro *and an adherent morphology of melanophores* in vivo

Given roles for JAM proteins in mediating cell–cell adhesion in other systems (Bazzoni, 2003; Ebnet, 2017), we asked whether zebrafish Jam3b promotes cell–cell adhesion and whether the missense substitution of *psr*^*vpr8e2*^ (A96E) lacks such activity. To these ends we quantified aggregation of *Drosophila* S2 cells transfected with cDNAs encoding wild-type or A96E mutant Jam3b following short term rotary culture (Eom et al., 2012). Cells expressing wild-type Jam3b formed large clusters within 60 min whereas cells expressing A96E mutant Jam3b or Actin-GFP (control) failed to aggregate in this manner (**Fig. 5A,B**). These results indicate the potential for Jam3b to promote cell adhesion during adult pigment pattern formation and the likelihood that A96E mutant Jam3b fails to support such activity *in vivo*.

**Fig. 5.**
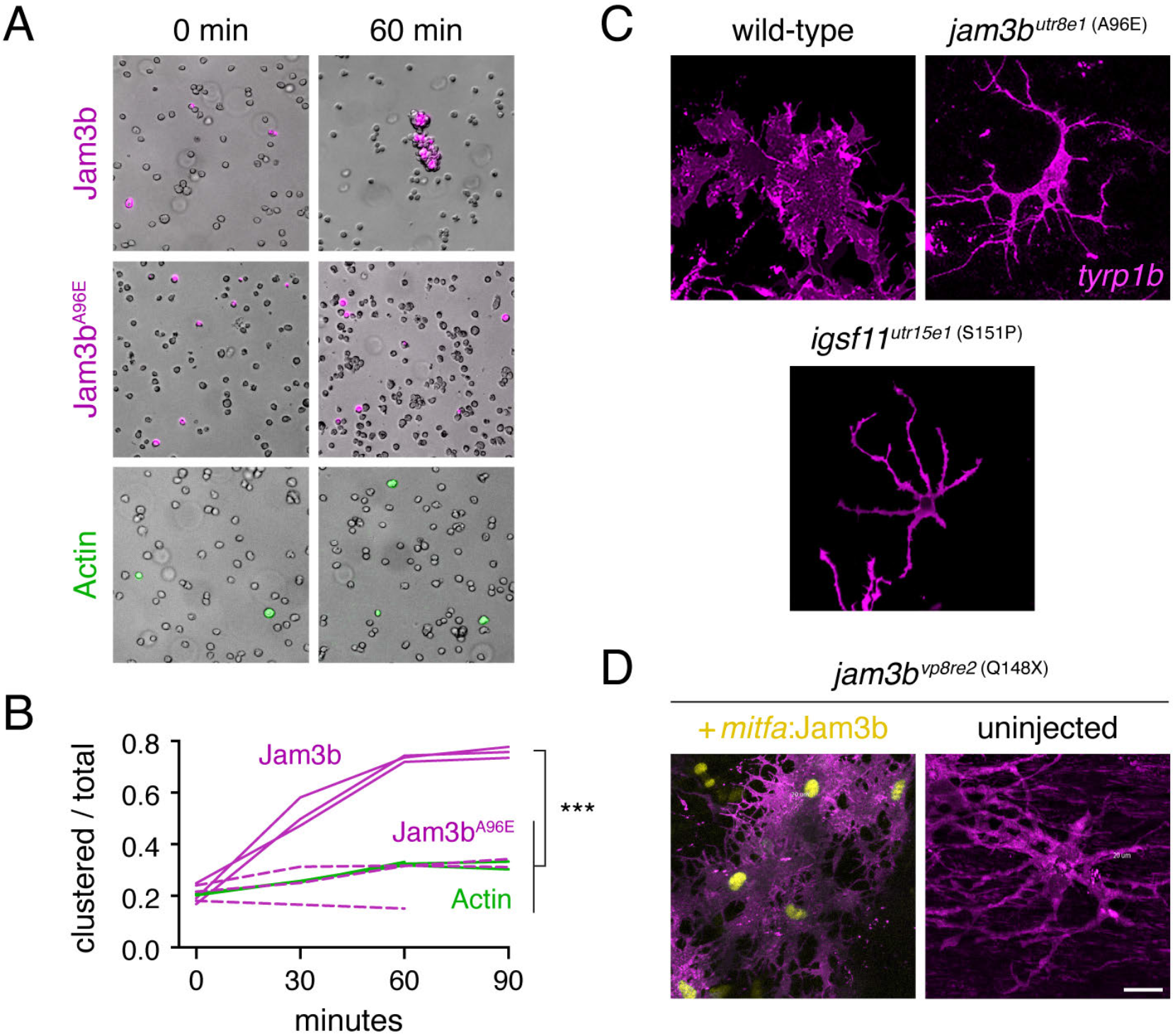
Jam3b promotes aggregation *in vitro* and an adherent morphology *in vivo*. (**A**) S2 cells transfected with constructs to express actin-GFP (control), wild-type Jam3b-mCherry or A96E mutant Jam3b-mCherry. Aggregates of cells expressing wild-type Jam3b formed within 60 minutes of rotary culture. (**B**) Quantification of transfected cells found within clusters relative to the total numbers of transfected cells in 3 replicate cultures. ***, Tukey-Kramer *post-hoc* comparisons of means *P*<0.0001 (overall ANOVA, *F*_2,6_=82.6, *P*<0.0001; data were arcsine-transformed for analysis to correct heterogeneity of variance among groups). (**C**) Membrane-targeted mCherry revealed well-spread morphologies of wild-type melanophores but more dendritic morphologies of *jam3b*^*utr8e1*^ and *igsf11*^*utr15e1*^ mutant melanophores *in vivo*. (**D**) Melanophores of *jam3b*^*vp8re2*^ mutants were also dendritic but adopted a more spread appearance when expressing wild-type Jam3b transgenically. Scale bars 20 *μ*M (D, for C and D).

If Jam3b normally promotes adhesion by melanophores, we predicted that melanophores of wild-type should be well-spread *in vivo* whereas melanophores of *jam3b* mutants should be more dendritic, typical of melanophores that are adherent, or compromised in their adhesive interactions, respectively (Eom et al., 2012; Milos et al., 1987). To delineate the outlines of melanophores we crossed TgBAC(*tyrp1b:PALM-mCherry*) (McMenamin et al., 2014) into mutant backgrounds and compared morphologies of cells expressing membrane-targeted mCherry between wild-type and *jam3b* mutants. Mature melanophores of wild-type fish were typically well-spread whereas melanophores of *jam3b* mutants were more often dendritic, and resembled those of *igsf11* mutant alleles (**Fig. 5C**), which also fail to support S2 cell aggregation (Eom et al., 2012). Defects in melanophore morphology in *jam3b* mutants were a direct consequence of Jam3b loss of function, as resupplying wild-type Jam3b transgenically rescued an adherent morphology (**Fig. 5D**) in addition to overall pattern (**Fig. 4D**).

### 3.6. Overlapping and non-overlapping requirements for Jam3b and Igsf11 during adult pigment pattern development

Repeated imaging, assays of cell autonomy and aggregation, and observations of melanophore morphologies suggested that Jam3b promotes adhesion and survival of melanophores, inferences similar to those for Igsf11 (Eom et al., 2012). To better understand the relative roles of these structurally similar adhesion receptors during pattern formation, we first asked whether Jam3b and Igsf11 are co-expressed in melanophores. Using a previously characterized polyclonal antiserum against zebrafish Igsf11 (Eom et al., 2012) we detected Igsf11 immunostaining in *jam3b*:Jam3b-mCherry+ melanophores *in vitro* (**Fig. 6A**).

**Fig. 6.**
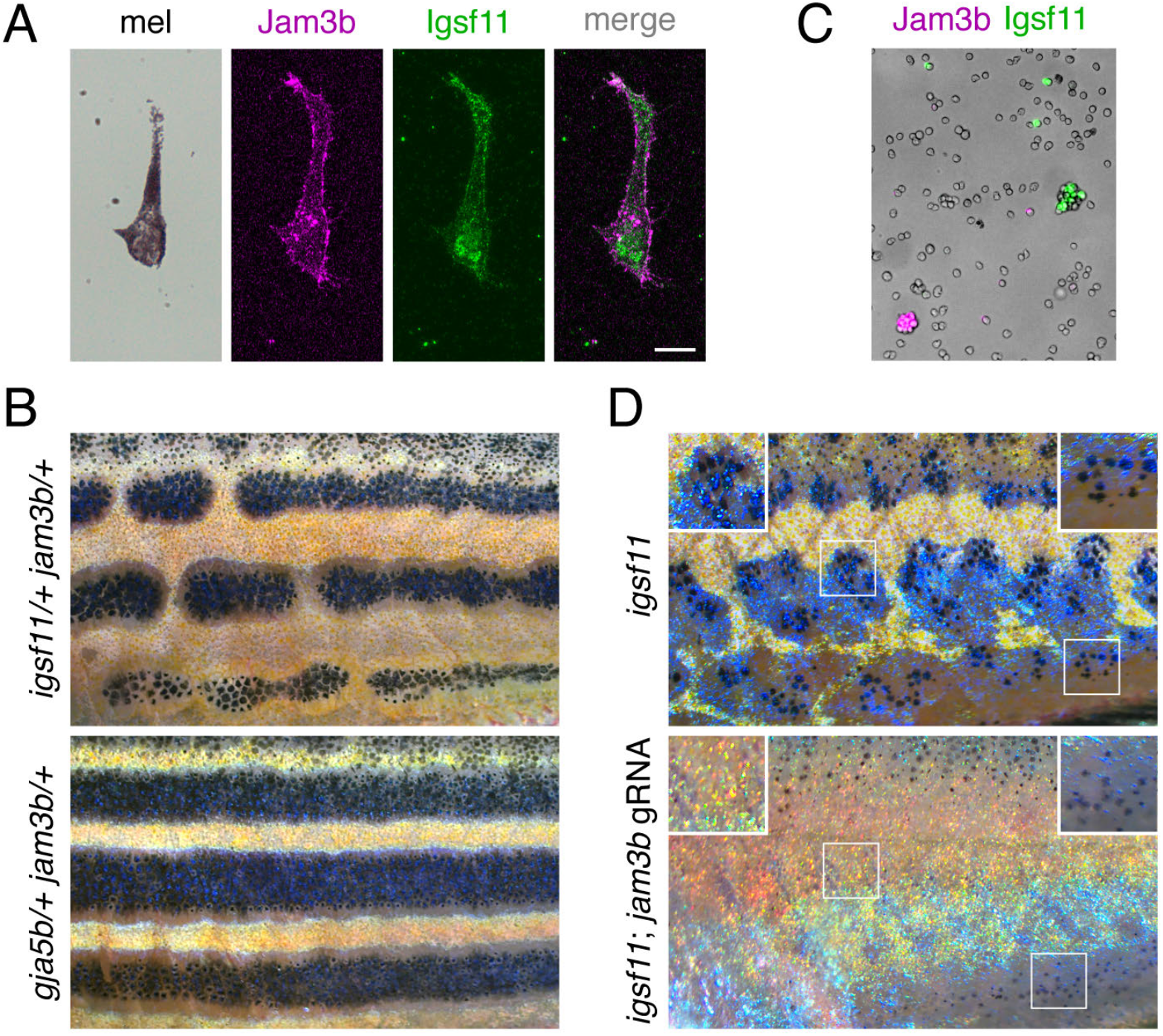
Expression and function of *jam3b* relative to *igsf11*. (**A**) Melanophores isolated from larvae (7.5 SSL) and plated onto culture dishes expressed *jam3b*:Jam3b-mCherry and stained with polyclonal antiserum to Igsf11. (**B**) Fish heterozygous for both *igsf11* and *jam3b* (here *igsf11*^*utr15e1*^ and *jam3b*^*vp8re2*^) had stripe breaks (*upper*) rather than wild-type stripes, as observed for single heterozygtoesheterozygous for fish heterozygous for both *gja5b* and *jam3b*^*vp8re2*^ (*lower*). (**C**) Jam3b-mCherry and Igsf11-EGFP supported homophilic but not heterophilic adhesion of S2 cells after 60 min in rotary culture. (**D**) Somatic loss of *jam3b* activity eliminated most residual hypodermal melanophores that develop in *igsf11* mutants (here *igsf11*^*vp15rc5*^). Insets show higher magnification views including some persisting melanophores at the ventral margin of the flank (*lower right*). Scale bar, 10 *μ*M (A).

Given overlap in expression for *igsf11* and *jam3b*, we interrogated overlap in function genetically. Mutant alleles of *jam3b* and *igsf11* are recessive, so we predicted that overlapping functions in melanophores might be revealed as epistatic phenotypes in fish doubly heterozyzous for both loci. Indeed, fish developed patterns with frequent stripe breaks when simultaneously heterozygous for missense allele *igsf11*^*wp15e1*^ (S151P) (Eom et al., 2012) and either *jam3b*^*utr8e1*^ (A96E) or *jam3b*^*vp8re2*^ (Q174X) (**Fig. 6B**, *upper*). To test if the phenotype was specific to this particular missense allele of *igsf11*, we generated a presumptive null allele, *igsf11*^*vp15rc4*^, having a 13 nucleotide insertion (M101in13) that leads to 15 novel amino acids and a premature stop codon in the first Ig-like domain. Fish heterozygous for *igsf11*^*vp15rc4*^ and *jam3b*^*vp8re2*^ developed the same pigment pattern defects, confirming the epistatic relationship between these loci (**Fig. S4A**). This genetic interaction was specific to *igsf11* and *jam3b*, as crosses of *jam3b*^*vp8re2*^ to the spotted *gja5b* (*leopard*) mutant, which resembles *igsf11* but encodes a gap junction protein, yielded fish with a wild-type pattern (**Fig. 6B**, *lower*). These results suggest that *jam3b* and *igsf11* have sufficiently overlapping functions that partial loss of function for either sensitizes pigment cells to partial loss of function for the other. This might, in principle, reflect a biochemical interaction between Jam3b and Igsf11. Nevertheless, we found that S2 cells separately expressing Jam3b or Igsf11 failed to aggregate in rotary culture (**Fig. 6C**) and Igsf11 immunostaining did not overlap with Jam3b-mCherry within melanophores (**Fig. 6A**), though we cannot exclude the possibility of fusion protein mislocalization, as noted above.

Because fish homozygous for either *jam3b* mutations or *igsf11* mutations retained substantial numbers of melanophores, we further asked whether residual melanophores represent a common population that develops independently of both loci. If so, fish doubly homozygous mutant for *jam3b* and *igsf11* should develop a pattern similar to either single mutant alone. To test this possibility, we induced somatic *jam3b* mutations in fish homozygous for a second presumptive null allele of *igsf11*, *igsf11*^*vp15rc5*^ (deletion of 8 nucleotides, M101del8, leading to a premature stop codon in the first Ig-like domain). Somatically double mutant fish had more severe phenotypes than either single mutant, lacking most hypodermal melanophores except for a population of lightly melanized cells near the ventral margin of the flank (**Fig. 6D**). We confirmed this phenotype by generating fish doubly homozygous for germline *jam3b*^*vp8re2*^ and *igsf11*^*vp15rc4*^ alleles, which also had variable numbers of residual melanophores ventrally (**Fig. S4B**). These results suggest that *jam3b* and *igsf11* have both overlapping and non-overlapping functions in hypodermal melanophore development.

Finally, we asked whether *jam3b* or *igsf11* are epistatic to another gene required for melanophore development, *kita*, encoding a Kit receptor tyrosine kinase. *kita* is required for survival and migration of embryonic melanophores, and promotes the establishment of latent progenitors to post-embryonic melanophores (Dooley et al., 2013; O’Reilly-Pol and Johnson, 2013; Parichy et al., 1999). In fish homozygous for null allele *kita*^*b5*^, only ~15% as many hypodermal melanophores differentiate as in wild-type, but those melanophores that do differentiate are abnormally proliferative, ultimately giving rise to a regulative population of melanophores with ~30% as many cells as in wild-type (Budi et al., 2011; Mills et al., 2007). Fish heterozygous for *jamb3b*^*utr8e1*^ and *kita*^*b5*^, or *igsf11*^*wp15e1*^ and *kita*^*b5*^, were phenotypically wild-type. By contrast, fish homozygous for *jamb3b*^*utr8e1*^ and *kita*^*b5*^ lacked all melanophores, and fish homozygous for missense allele *igsf11*^*utr15e1*^ and *kita*^*b5*^ lacked nearly all melanophores (**Fig. S4C**). These phenotypes show that both *jam3b* and *igsf11* are required by regulative melanophores that develop in the absence of *kita* activity.

## 4. Discussion

The repertoire of genes essential for adhesive interactions by cells, and how individual genes and higher order interactions help to choreograph morphogenesis and differentiation in developing organisms remain largely mysterious. We have shown that the two Ig-domain adhesion receptor Jam3b is required by melanophores, and for the adult pigment pattern to which these cells contribute. Jam3b allowed the aggregation of heterologous cells *in vitro* and supported survival and an adherent phenotype of melanophores *in vivo*, activities partly but not completely overlapping with Igsf11 and Kita. Our study provides a new entry point for dissecting roles of adhesion in melanophore development and hints at the complexity of adhesive interactions during pattern development.

A parsimonious model of *jam3b* function in pigment pattern formation would have an initial requirement in promoting melanophore survival during the larva-to-adult transformation, when deficits in melanophore numbers and loss of melanophores were first apparent in *jamb3* mutants (**Fig. 1C,D,F**). A role in melanophore survival during later homeostasis would be present too, as adult *jam3b* mutants exhibited melanized debris and gaps in residual stripes, where bluish iridophores occurred without melanophores (**Fig. S2B,C**), despite these iridophores normally associating with melanophores and requiring them for their development (Gur et al., 2020). In this model, Jam3b acts autonomously to melanophores to promote their adhesion, as evidenced by cell transplantation and transgene rescue. Though *jam3b* is transcribed in pigment cell progenitors (**Fig. 3B**), it would become essential in the melanophore lineage only after melanization is underway, a requirement extending through terminal maturation, when a spread morphology has been acquired (**Fig. 5C,D**) and when Jam3b-mCherry was seen to have accumulated at sites of overlap between melanophores in the stripe (**Fig. 3E**). The cellular consequences of Jam3b activity would be to promote adhesive contacts with other cells required for melanophore survival, allowing a cohesive, stable phenotype of the stripes in which melanophores are arranged. Lacking such activity, melanophores fail to survive and stripes fail to acquire or maintain their normal integrity.

JAM3 proteins of other species promote adhesive interactions between cells, assembling at cell–cell contacts including adherens junctions and tight junctions and interacting with Zonula Occludens-1 (ZO-1) and ZO-2; interactions with other scaffolding proteins are likely, as are roles in assembling signaling complexes where cells contact one another (Aurrand-Lions et al., 2001; Ebnet, 2017; Ebnet et al., 2003; Economopoulou et al., 2009; Morris et al., 2006; Reymond et al., 2005; Satohisa et al., 2005). In zebrafish melanophores, these activities could occur in either of two broad contexts, each having different implications for pattern formation.

In one scenario, Jam3b functions specifically in melanophores to promote interactions with other melanophores, likely through homophilic interactions in *trans* with Jam3b, or heterophilic interactions in *trans* with Jam2a as observed in myoblasts (Powell and Wright, 2011). This is an attractive idea, given localization of Jam3b-mCherry at sites of overlap between melanophore membranes. Indeed, stable contacts are made between melanophores of another species, the newt *Taricha torosa*, during a terminal consolidation of larval stripes (Parichy, 1996; Twitty, 1945). A similar process could occur in zebrafish and such interactions might be reflected in a “community effect” on melanophore patterning, in which a threshold cell density would be required to support melanophore survival and stripe integrity. The development of more severe patterns in individually reared and quickly growing *jam3b* mutants is consistent with melanophores failing to achieve such a critical density in a rapidly expanding tissue environment. A similar phenomenon is seen in the alpha tubulin mutant *puma* (*tuba8l3*), which exhibits more severe defects when reared in high somatic growth conditions than in low somatic growth conditions (Larson et al., 2010; Parichy and Turner, 2003b). In both *jam3b* and *puma* mutants, patterns are highly variegated and differ in severity from individual to individual, features that are qualitatively different from many other pattern mutants. Community effects are also consistent with the death of melanophores that become isolated after failing to join stripes during normal development (Parichy and Turner, 2003b; Takahashi and Kondo, 2008), and have been invoked to explain the variegated melanocyte phenotype of *patchwork* mutant mice (Aubin-Houzelstein et al., 1998).

If Jam3b does promote melanophore–melanophore adhesion and trophic support, it likely does so through interactions that support contact in a permissive environment, but not a true adhesive tethering of melanophores to one another: if xanthophores are removed from interstripes, melanophores readily disperse from stripes, showing they are free to move when there is opportunity to do so (Parichy and Turner, 2003a; Takahashi and Kondo, 2008). Our observation that Jam3b-mCherry accumulated in membranes where melanophores overlapped, but was largely absent from regions of membrane without adjacent melanophores at the stripe– interstripe border, further raises the possibility that Jam3b helps to impart a polarity to stripe melanophores. Indeed, JAM3 is required for the establishment of polarity during spermatogenesis through interactions with Partitioning-defective (Par) proteins (Gliki et al., 2004) and may have similar roles in endothelial development (Ebnet, 2017; Ebnet et al., 2003).

In a second, complementary scenario, Jam3b functions in melanophores but via interactions with other types of cells in the local environment. For example, there would seem to be ample opportunity for Jam3b-dependent interactions with iridophores: interstripe iridophores influence melanophore localization (Frohnhofer et al., 2013; Patterson et al., 2014; Patterson and Parichy, 2013), stripe iridophores require melanophores for their development (Gur et al., 2020), and iridophores express both *jam3b* and *jam2a*. Indeed, one interpretation of our rescue experiment—in which mutant melanophores that expressed Jam3b transgenically recovered an adherent phenotype and formed normal stripes—would be that Jam3b allowed melanophores to interact with Jam2a, or another JAM protein, expressed by nearby iridophores. Although we cannot exclude this possibility we do not presently favor it. Were this a major function of Jam3b and a major contributor to melanophore–iridophore interactions we would have expected fish lacking iridophores to have a similar melanophore phenotype regardless of Jam3b activity. Yet melanophore phenotypes of iridophore-free fish depended on whether they were wild-type or mutant for *jam3b*, and *jam3b* mutants had similar phenotypes whether or not they had iridophores. Assessing additional roles for Jam3b-dependent interactions with iridophores, xanthophores, or other hypodermal cells (Aman and Parichy, 2020), will require additional genetic, and genetic mosaic analyses, and could yield interesting insights. Indeed, an emerging view is that JAM proteins also have major roles in regulating motility and signaling via interactions in *cis* with integrins (Kummer and Ebnet, 2018; Mandicourt et al., 2007), and roles for regional extracellular matrices in zebrafish pigment pattern development remain wholly unexplored.

Beyond roles autonomous to hypodermal melanophores, Jam3b may also function in iridophores, in which transcripts and Jam3b-mCherry expression were detected. Though iridophore development in the absence of *jam3b* was overtly normal for a background deficient in melanophores (Frohnhofer et al., 2013; Gur et al., 2020), the accumulation of Jam3b-mCherry where interstripe iridophores were juxtaposed with one another suggests potential roles in establishing or maintaining the dense epithelial-like arrangement of these cells. Normal iridophore patterning requires ZO-1 (encoded in zebrafish by *tight junction protein 1a*) (Fadeev et al., 2015) and zebrafish ZO-1 is likely to interact with Jam3b, as ZO-1 of other species can interact with other JAM family members (Ebnet, 2017; Steinbacher et al., 2018). These and other functions of Jam3b in pigment cell lineages are currently under investigation.

Jam3b is the second 2-Ig domain adhesion receptor implicated in melanophore patterning and our analyses allow for an initial parsing of its roles relative to the first, Igsf11, and also the better understood receptor tyrosine kinase, Kita. In zebrafish, Igsf11 promotes migration and survival of melanophores and melanoblasts (Eom et al., 2012). In *jam3b* mutants, behaviors of melanoblasts were similar to wild-type, suggesting that the requirement for Jam3b arises somewhat later than that for Igsf11 in this lineage. JAM and CAR receptors can interact with the same partners in other systems (Ebnet, 2017; Rathjen, 2020), and a genetic interaction— revealed as broken stripes in doubly heterozygous fish—suggests some functional overlap in zebrafish melanophores. Indeed, melanophores mutant for either gene had more dendritic phenotypes. It thus seems likely that products of both genes contribute to contacts between melanophores or other cells essential for signaling, survival and an adherent morphology, and that partially reduced activity at either locus can normally be compensated by the other locus. Compensatory mechanisms between JAM, JAM-like and CAR proteins have been inferred in other system as well (Ebnet, 2017).

Our finding that homozygous *jam3b; igsf11* mutants lacked nearly all melanophores suggests that Jam3b and Igsf11 have non-overlapping roles as well: perhaps Jam3b functions principally at adherens or tight junctions, whereas Igsf11 functions principally at gap junctions (Rathjen, 2020), which play essential roles in zebrafish adult pattern development (Irion et al., 2014; Usui et al., 2019; Watanabe et al., 2006; Watanabe et al., 2016). Interestingly, the melanophores that were found in double mutants were lightly melanized and limited to ventral regions of the flank, where some melanoblasts emerge during the larva-to-adult transition (Budi et al., 2011). This phenotype hints at additional roles for Jam3b in motility and differentiation, revealed only in the absence of Igsf11. Partially non-overlapping roles for Jam3b and Kita were likewise suggested by the loss of all melanophores in fish lacking both receptors. Jam3b may have an especially critical role in allowing adhesion and survival of the few, large and proliferative melanophores that develop without Kita signaling.

Finally, our identification of *jam3b* requirements in development and patterning of zebrafish melanophores may have implications for studies of melanoma. Functions of JAM3 in human melanocyte development have not been documented. Yet human and mouse melanoma cells can express JAM3, and higher levels of expression are associated with interactions with JAM2 expressed by endothelial cells and enhanced extravasation from the vasculature, allowing metastasis to peripheral locations including the lung (Arcangeli et al., 2012; Ghislin et al., 2011; Langer et al., 2011). Zebrafish has become a valuable model for studying the initiation and subsequent metastasis of melanoma cells (Kaufman et al., 2016; Kim et al., 2017; Neiswender et al., 2017; Patton et al., 2021; Roh-Johnson et al., 2017) and *jam3b* mutants could be useful in further dissecting these processes.

## Acknowledgements

For assistance we thank Michael Lang (positional cloning), Jessica Turner, Marianne Cole, Amber Schwindling, Samir Halabiya and Tiffany Gordon (fish rearing and non-complementation screening), Deqwon Pendergrass (genotyping), Emily Bain (imaging and chimeras), Andy Aman (whole fish Jam3-mCherry imaging), and Braedan McCluskey (melanophore counts). Supported by NIH R35 GM122471 to DMP and NIH T32 GM008136 to RRB.

---

**Fig. S1.**
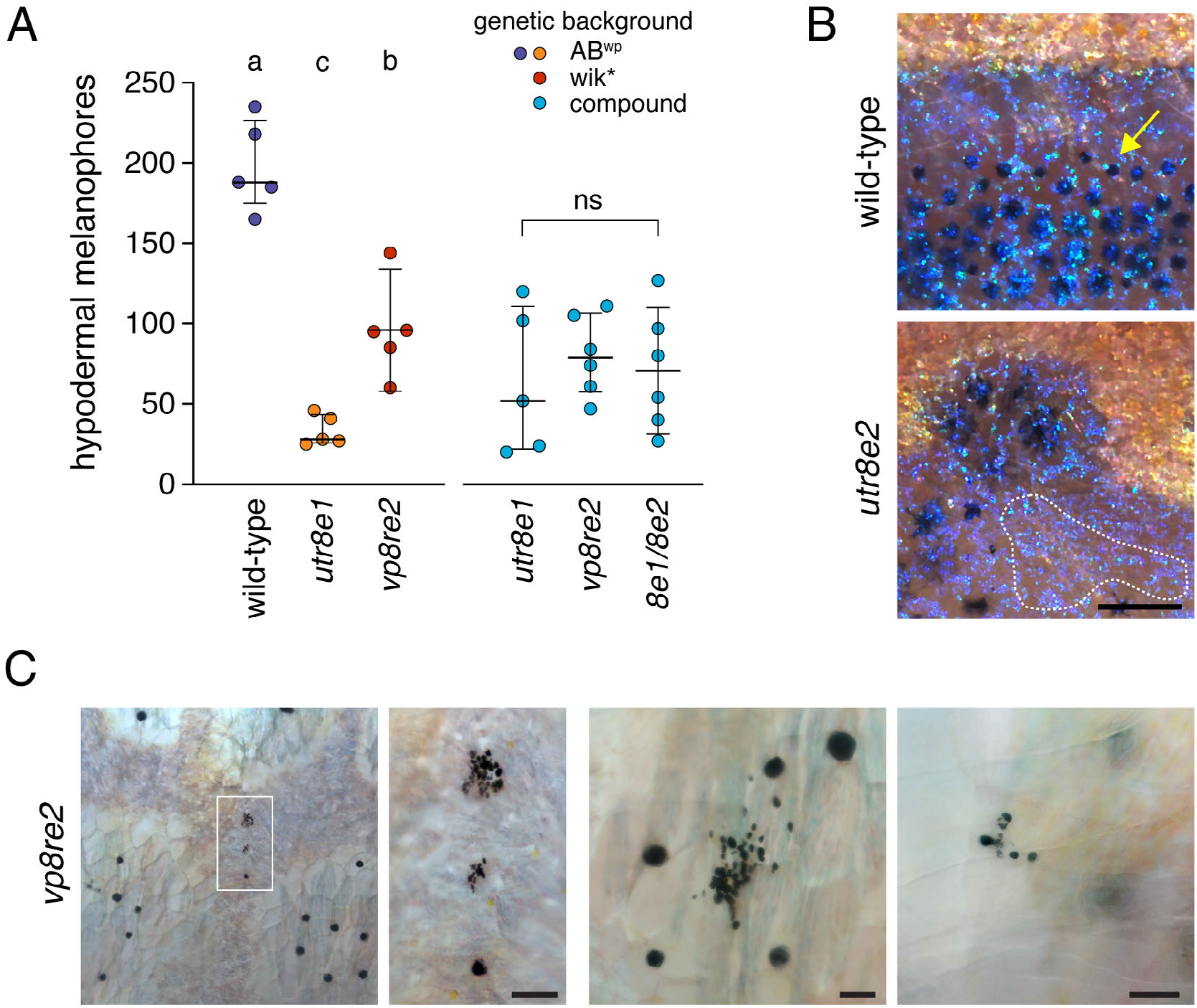
Pigmentary phenotypes of *jam3b* mutant adults. (**A**) Numbers of hypodermal melanophores along five segments at the level of the ventral primary melanophore stripe of wild-type and dorsal to the anal fin. Left side of plot indicates melanophores in wild-type and homozygous *psr*^*utr8e1*^ or *psr*^*vp8re2*^ mutants, as maintained in different genetic backgrounds. Each point represents an individual fish and bars show means ± 95% confidence intervals. Different letters indicate means significantly different in *post hoc* Tukey-Kramer comparisons (all *P*<0.005; overall ANOVA: *F*_2,12_=57.6, *P*<0.0001). Right side of plot shows lack of significant differences among allelic combinations in a common genetic background (*F*_2,14_=0.3, *P*=0.7488). (**B**) Details of iridophores typical of interstripes (yellow-gold iridescent cells) and stripes (blue iridescent cells) in both wild-type and *psr* mutants. Although blue stripe iridophores normally require melanophores for their development (Gur et al., 2020), patches of such iridophores that lacked associated melanophores were often evident in *psr* mutants (dashed line). Note that melanophore cell bodies and processes extend beyond regions of visible melanin (e.g., arrow) (Hamada et al., 2014), which concentrates towards cell centers during normal physiological modulation of melanosome dispersion and when fish are treated with epinephrine to contract melanosomes from the periphery (Fujii, 2000). Yellow–orange xanthophores are faintly visible over interstripe iridophores. (**C**) Melanized cell fragments in adult *psr*^*vp8re2*^ mutants. Left two panels, low magnification and detail of melanized debris in hypodermis. Right panels show additional examples in hypodermis and at the surface of a scale, typical of debris being extruded through the epidermis (Lang et al., 2009). Scale bars, 200 *μ*M (B), 50 *μ*M (C).

**Fig. S2.**
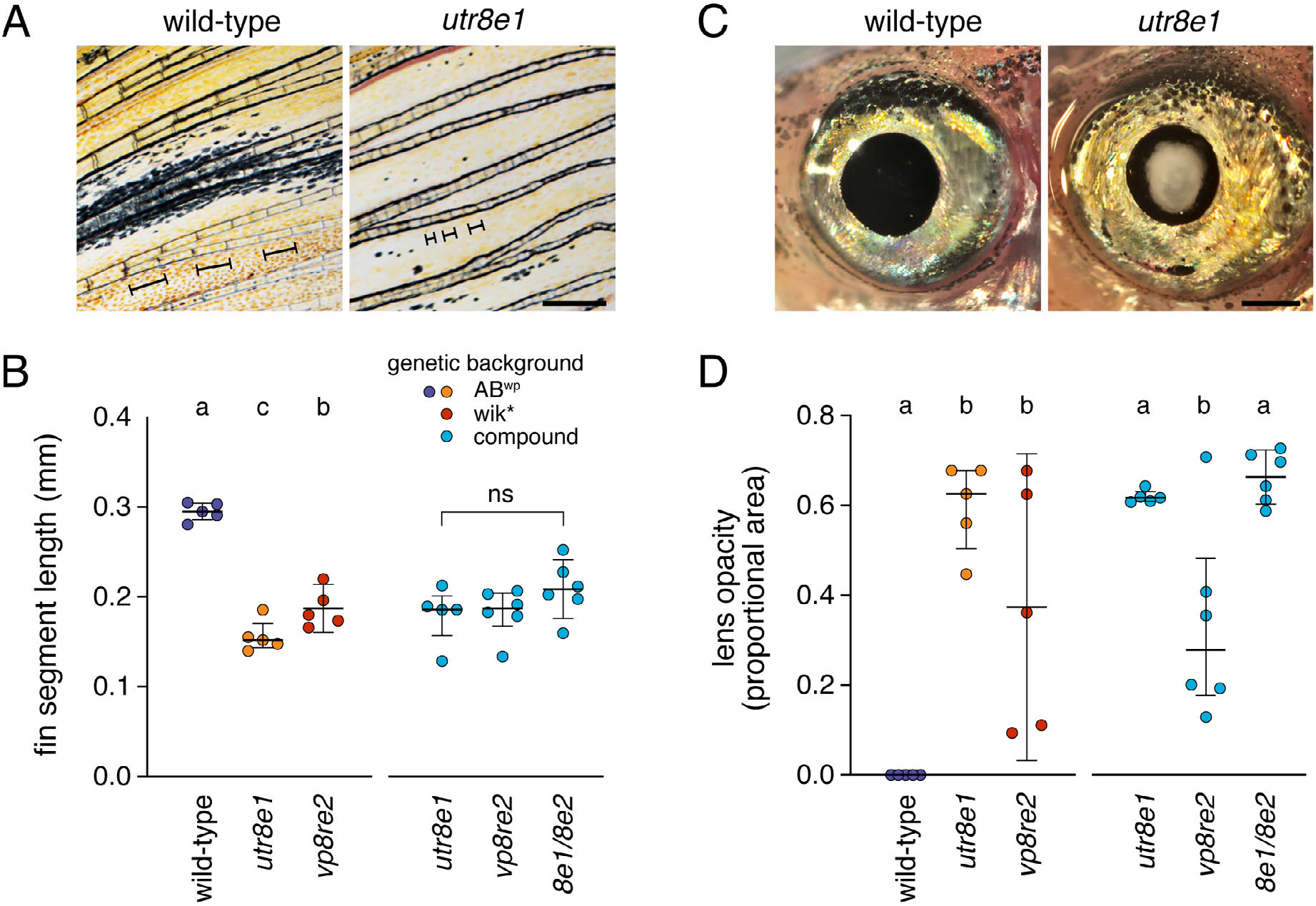
Fin and lens phenotypes of *psr* mutants. (**A**) Lepidotrichia of *psr* mutants were shorter and more variable in morphology than wild-type. Brackets in images indicate lengths of individual lepidotrichia. (**B**) Though alleles differed in lepidotrichial length when maintained in separate genetic backgrounds, combinations of mutant alleles did not differ in a common genetic background, similar to results for melanophores. Left side, Tukey-Kramer comparisons all *P*<0.05; overall *F*_2,12_=92.6, *P*<0.0001. Right side, overall *F*_2,14_=1.6, *P*=0.2350. (**C**) Lens opacity in *psr*^*utr8e1*^ adult at ~1 year. (**D**) Proportional areas of opacities differed between alleles in original backgrounds and also in a common background. Left side, overall *F*_2,12_=14.8, *P*<0.0001. Right side, overall *F*_2,14_=10.7, *P*=0.0015. Tukey-Kramer comparisons of means all *P*<0.05 when letters above bars differ. Scale bars, 500 *μ*M.

**Fig. S3.**
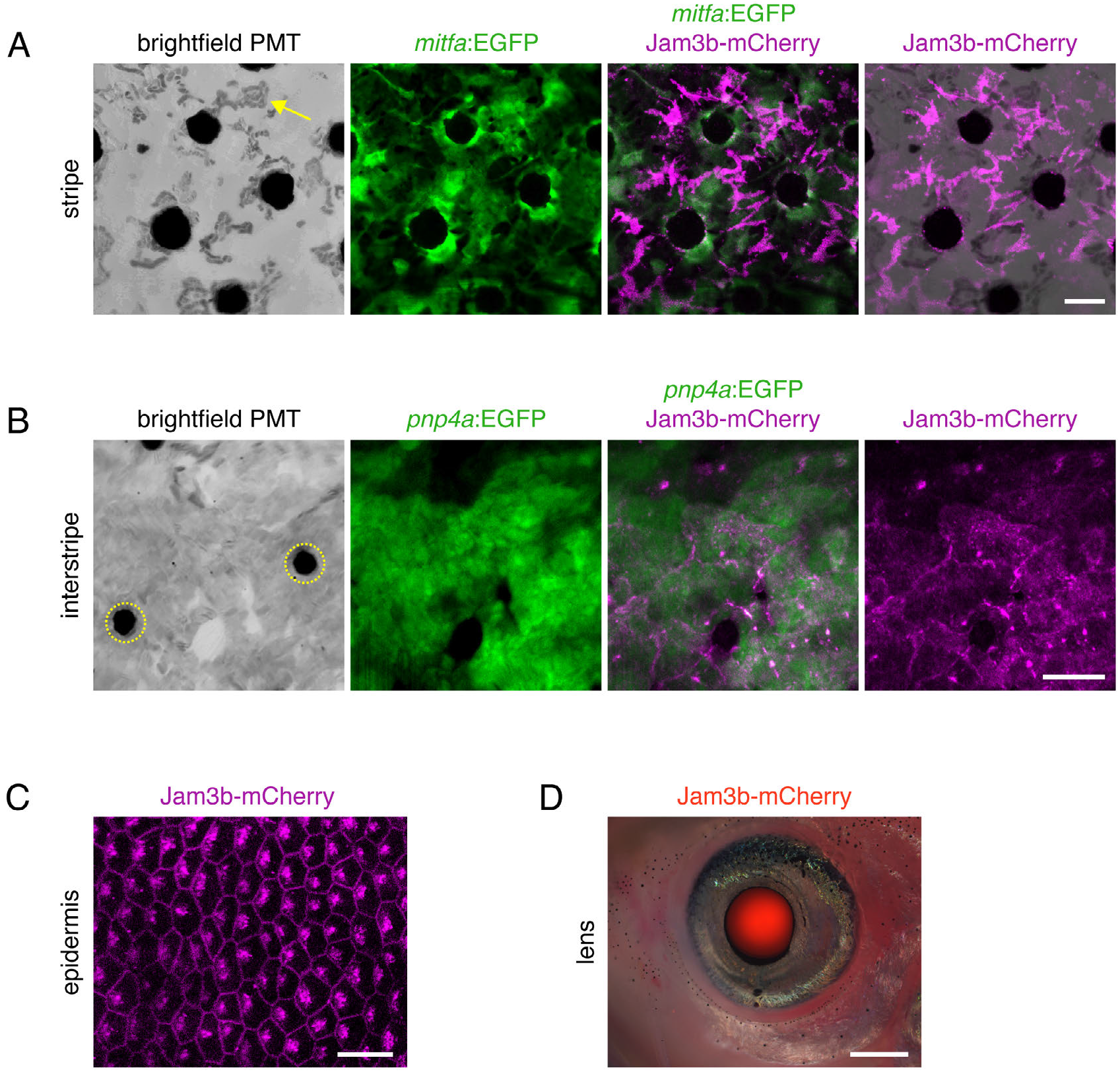
*jam3b*:Jam3b-mCherry expression during post-embryonic development. (**A**) Melanophores of the stripe. In brightfield melanin is evident, contracted by epinephrine to centers of melanophores. Reflecting platelets of sparsely arranged stripe iridophores are also evident (e.g., arrow). Melanophores express *mitfa*:EGFP, with *jam3b*:Jam3b-mCherry accumulations at cell peripheries. Occasional melanophores within stripes lacked pronounced Jam3b-mCherry expression (lower edge of frame), perhaps owing to transgene variegation or shorter times since differentiation. In contrast to melanophores beneath them, stripe iridophores did not have detectable accumulation of Jam3b-mCherry. (**B**) In the interstripe, reflecting platelets of densely packed iridophores are apparent. In brightfield captured by photomultiplier tube (PMT), accumulations of carotenoid pigments within xanthophores present in a place aboce iridophores appear black (dashed circles). Iridophores and reflecting platelets are labeled within *pnp4a*:mem-mEGFP and Jam3b-mCherry is evident where cells contact one another. (**C**) Basal epidermal cells of the skin. (**D**) Superimposed brightfield and fluorescence of image of Jam3b-mCherry expression in lens of a jvenile fish. Scale bars, 20 *μ*M (A,B,C) 500 *μ*M (D).

**Fig. S4.**
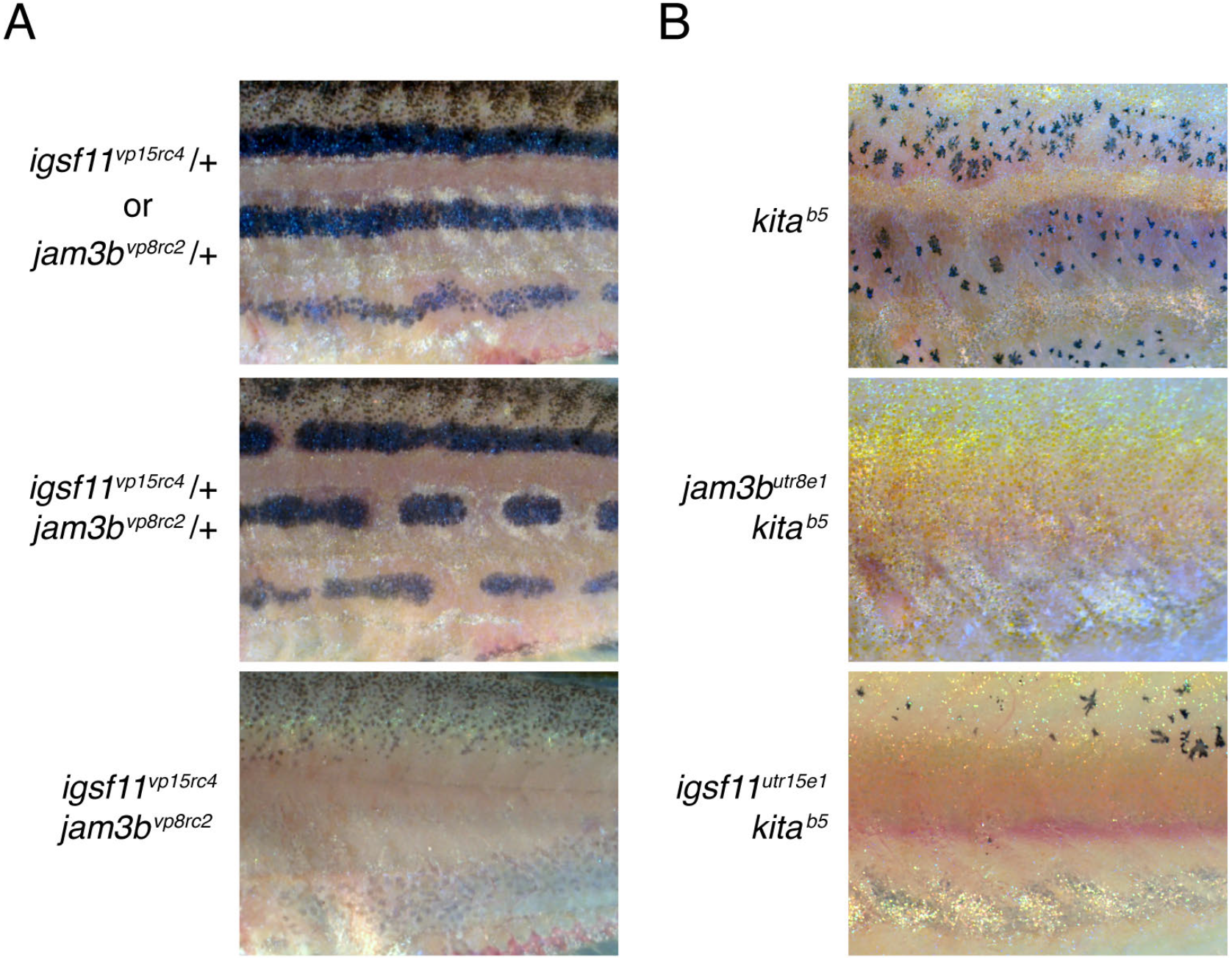
*jam3b* dependent melanophore development in mutants of *igsf11* and *kita*. (**A**) Fish heterozygous for *igsf11*^*vp15rc4*^ or *jam3b*^*vp8rc2*^ had pigment patterns with only occasional breaks in otherwise wild-type stripes (upper). Fish heterozygous for both mutations had numerous stripe breaks (middle) and fish homozygous for both mutations lacked hypodermal melanophores over the middle flank but often developed lightly melanized melanophores along the ventral margin of the flank (lower). (**B**) A population of regulative melanophores develops in homozygous *kita* mutants (upper), but these cells were entirely missing (*middle*), or further reduced in number (*lower*), by simultaneous homozygous mutation of *jam3b* or *igsf11*.

**Movie S1. Development of pigment pattern in wild-type.** Melanophore pattern emerges gradually as these cells differentiate and stripes become increasingly well organized. Images shown were compiled for a single representative individual, taken on successive days and rescaled to control for growth (corresponding to still image details in **Fig. 1C**). Imaging was terminated before adult patterning had been completed and while some melanophores still persist in the interstripe. Differences in melanophore morphology across days represent physiological variation in melanosome dispersion within cells.

**Movie S2. Development of pigment pattern in***psr* **mutant.** In contrast to wild-type, melanophores were more likely to disappear from the pattern, indicative of cell death, and persisting melanophores tended to have smaller melanized areas. As melanophores were lost, yellowish, densely packed iridophores typical of the interstripe came to populate regions of the flank that would normally be occupied by melanophores and bluish iridophores of the stripes.

**Movie S3. *mitfa*:GFP+ melanoblasts of wild-type.** Motile spindle shaped cells are melanoblasts that will differentiate as melanophores (Budi et al., 2011). Shown is a region of the ventral flank in an explant at 7 SSL. Some persisting early larval melanophores are evident along the horizontal myoseptum near the top of the frame. Scattered non-motile fluorescent puncta are accumulations of autofluorescent carotenoids in developing xanthophores, appearance of which varies somewhat from indvidual to individual. Movie shown is representative of >10 trunks examined. Frame interval, 30 min.

**Movie S4. *mitfa*:GFP+ melanoblasts of *psr* mutant.** Cell retained motile activity and did not exhibit an overt failure of survival as observed in some mutants (Budi et al., 2011). Movie shown is representative of >10 trunks examined. Frame interval, 30 min.

## Notes

### Competing Interest Statement

The authors have declared no competing interest.

## References

Adameyko, I., Lallemend, F., Aquino, J.B., Pereira, J.A., Topilko, P., Muller, T., Fritz, N., Beljajeva, A., Mochii, M., Liste, I., Usoskin, D., Suter, U., Birchmeier, C., Ernfors, P., 2009. Schwann cell precursors from nerve innervation are a cellular origin of melanocytes in skin. Cell 139, 366–379.

Aman, A.J., Parichy, D.M., 2020. Chapter 8 - Zebrafish Integumentary System, in: Cartner, S.C., Eisen, J.S., Farmer, S.C., Guillemin, K.J., Kent, M.L., Sanders, G.E. (Eds.), The Zebrafish in Biomedical Research. Academic Press, pp. 91–96.

Arcangeli, M.L., Frontera, V., Bardin, F., Thomassin, J., Chetaille, B., Adams, S., Adams, R.H., Aurrand-Lions, M., 2012. The Junctional Adhesion Molecule-B regulates JAM-C-dependent melanoma cell metastasis. FEBS Lett 586, 4046–4051.

Aubin-Houzelstein, G., Bernex, F., Elbaz, C., Panthier, J.J., 1998. Survival of patchwork melanoblasts is dependent upon their number in the hair follicle at the end of embryogenesis. Dev Biol 198, 266–276.

Aurrand-Lions, M., Johnson-Leger, C., Wong, C., Du Pasquier, L., Imhof, B.A., 2001. Heterogeneity of endothelial junctions is reflected by differential expression and specific subcellular localization of the three JAM family members. Blood 98, 3699–3707.

Bazzoni, G., 2003. The JAM family of junctional adhesion molecules. Curr Opin Cell Biol 15, 525–530.

Budi, E.H., Patterson, L.B., Parichy, D.M., 2011. Post-embryonic nerve-associated precursors to adult pigment cells: genetic requirements and dynamics of morphogenesis and differentiation. PLoS Genet 7, e1002044.

Cao, J., Spielmann, M., Qiu, X., Huang, X., Ibrahim, D.M., Hill, A.J., Zhang, F., Mundlos, S., Christiansen, L., Steemers, F.J., Trapnell, C., Shendure, J., 2019. The single-cell transcriptional landscape of mammalian organogenesis. Nature 566, 496–502.

Choi, J., Dong, L., Ahn, J., Dao, D., Hammerschmidt, M., Chen, J.N., 2007. FoxH1 negatively modulates flk1 gene expression and vascular formation in zebrafish. Dev Biol 304, 735–744.

Curran, K., Lister, J.A., Kunkel, G.R., Prendergast, A., Parichy, D.M., Raible, D.W., 2010. Interplay between Foxd3 and Mitf regulates cell fate plasticity in the zebrafish neural crest. Dev Biol 344, 107–118.

Dilshat, R., Fock, V., Kenny, C., Gerritsen, I., Lasseur, R.M.J., Travnickova, J., Eichhoff, O.M., Cerny, P., Moller, K., Sigurbjornsdottir, S., Kirty, K., Einarsdottir, B.O., Cheng, P.F., Levesque, M., Cornell, R.A., Patton, E.E., Larue, L., de Tayrac, M., Magnusdottir, E., Helga Ogmundsdottir, M., Steingrimsson, E., 2021. MITF reprograms the extracellular matrix and focal adhesion in melanoma. 10.

Dooley, C.M., Mongera, A., Walderich, B., Nusslein-Volhard, C., 2013. On the embryonic origin of adult melanophores: the role of ErbB and Kit signalling in establishing melanophore stem cells in zebrafish. Development 140, 1003–1013.

Ebnet, K., 2017. Junctional Adhesion Molecules (JAMs): Cell Adhesion Receptors With Pleiotropic Functions in Cell Physiology and Development. Physiol Rev 97, 1529–1554.

Ebnet, K., Aurrand-Lions, M., Kuhn, A., Kiefer, F., Butz, S., Zander, K., Meyer zu Brickwedde, M.K., Suzuki, A., Imhof, B.A., Vestweber, D., 2003. The junctional adhesion molecule (JAM) family members JAM-2 and JAM-3 associate with the cell polarity protein PAR-3: a possible role for JAMs in endothelial cell polarity. J Cell Sci 116, 3879–3891.

Ebnet, K., Suzuki, A., Ohno, S., Vestweber, D., 2004. Junctional adhesion molecules (JAMs): more molecules with dual functions? J Cell Sci 117, 19–29.

Economopoulou, M., Hammer, J., Wang, F., Fariss, R., Maminishkis, A., Miller, S.S., 2009. Expression, localization, and function of junctional adhesion molecule-C (JAM-C) in human retinal pigment epithelium. Invest Ophthalmol Vis Sci 50, 1454–1463.

Eom, D.S., Inoue, S., Patterson, L.B., Gordon, T.N., Slingwine, R., Kondo, S., Watanabe, M., Parichy, D.M., 2012. Melanophore migration and survival during zebrafish adult pigment stripe development require the immunoglobulin superfamily adhesion molecule Igsf11. PLoS Genet 8, e1002899.

Erickson, C.A., Perris, R., 1993. The role of cell-cell and cell-matrix interactions in the morphogenesis of the neural crest. Dev Biol 159, 60–74.

Fadeev, A., Krauss, J., Frohnhofer, H.G., Irion, U., Nusslein-Volhard, C., 2015. Tight Junction Protein 1a regulates pigment cell organisation during zebrafish colour patterning. eLife 4.

Frohnhofer, H.G., Krauss, J., Maischein, H.M., Nusslein-Volhard, C., 2013. Iridophores and their interactions with other chromatophores are required for stripe formation in zebrafish. Development 140, 2997–3007.

Fujii, R., 2000. The regulation of motile activity in fish chromatophores. Pigment Cell Res 13, 300–319.

Garcia-Lopez, M.A., Barreiro, O., Garcia-Diez, A., Sanchez-Madrid, F., Penas, P.F., 2005. Role of tetraspanins CD9 and CD151 in primary melanocyte motility. J Invest Dermatol 125, 1001–1009.

Ghislin, S., Obino, D., Middendorp, S., Boggetto, N., Alcaide-Loridan, C., Deshayes, F., 2011. Junctional adhesion molecules are required for melanoma cell lines transendothelial migration in vitro. Pigment Cell Melanoma Res 24, 504–511.

Gliki, G., Ebnet, K., Aurrand-Lions, M., Imhof, B.A., Adams, R.H., 2004. Spermatid differentiation requires the assembly of a cell polarity complex downstream of junctional adhesion molecule-C. Nature 431, 320–324.

Gur, D., Bain, E.J., Johnson, K.R., Aman, A.J., Pasoili, H.A., Flynn, J.D., Allen, M.C., Deheyn, D.D., Lee, J.C., Lippincott-Schwartz, J., Parichy, D.M., 2020. In situ differentiation of iridophore crystallotypes underlies zebrafish stripe patterning. Nat Commun 11, 6391.

Haage, A., Wagner, K., Deng, W., Venkatesh, B., Mitchell, C., Goodwin, K., Bogutz, A., Lefebvre, L., Van Raamsdonk, C.D., Tanentzapf, G., 2020. Precise coordination of cell-ECM adhesion is essential for efficient melanoblast migration during development. Development 147.

Haass, N.K., Smalley, K.S., Li, L., Herlyn, M., 2005. Adhesion, migration and communication in melanocytes and melanoma. Pigment Cell Res 18, 150–159.

Hamada, H., Watanabe, M., Lau, H.E., Nishida, T., Hasegawa, T., Parichy, D.M., Kondo, S., 2014. Involvement of Delta/Notch signaling in zebrafish adult pigment stripe patterning. Development 141, 318–324.

Hirata, M., Nakamura, K., Kanemaru, T., Shibata, Y., Kondo, S., 2003. Pigment cell organization in the hypodermis of zebrafish. Dev Dyn 227, 497–503.

Inaba, M., Yamanaka, H., Kondo, S., 2012. Pigment pattern formation by contact-dependent depolarization. Science 335, 677.

Irion, U., Frohnhofer, H.G., Krauss, J., Colak Champollion, T., Maischein, H.M., Geiger-Rudolph, S., Weiler, C., Nusslein-Volhard, C., 2014. Gap junctions composed of connexins 41.8 and 39.4 are essential for colour pattern formation in zebrafish. eLife 3, e05125.

Johnson, S.L., Africa, D., Horne, S., Postlethwait, J.H., 1995. Half-tetrad analysis in zebrafish: mapping the ros mutation and the centromere of linkage group I. Genetics 139, 1727–1735.

Kaufman, C.K., Mosimann, C., Fan, Z.P., Yang, S., Thomas, A.J., Ablain, J., Tan, J.L., Fogley, R.D., van Rooijen, E., Hagedorn, E.J., Ciarlo, C., White, R.M., Matos, D.A., Puller, A.C., Santoriello, C., Liao, E.C., Young, R.A., Zon, L.I., 2016. A zebrafish melanoma model reveals emergence of neural crest identity during melanoma initiation. Science 351, aad2197.

Kelsh, R.N., Harris, M.L., Colanesi, S., Erickson, C.A., 2009. Stripes and belly-spots -- a review of pigment cell morphogenesis in vertebrates. Semin Cell Dev Biol 20, 90–104.

Kim, I.S., Heilmann, S., Kansler, E.R., Zhang, Y., Zimmer, M., Ratnakumar, K., Bowman, R.L., Simon-Vermot, T., Fennell, M., Garippa, R., Lu, L., Lee, W., Hollmann, T., Xavier, J.B., White, R.M., 2017. Microenvironment-derived factors driving metastatic plasticity in melanoma. Nat Commun 8, 14343.

Kobayashi, I., Kobayashi-Sun, J., Hirakawa, Y., Ouchi, M., Yasuda, K., Kamei, H., Fukuhara, S., Yamaguchi, M., 2020. Dual role of Jam3b in early hematopoietic and vascular development. Development 147.

Kostrewa, D., Brockhaus, M., D’Arcy, A., Dale, G.E., Nelboeck, P., Schmid, G., Mueller, F., Bazzoni, G., Dejana, E., Bartfai, T., Winkler, F.K., Hennig, M., 2001. X-ray structure of junctional adhesion molecule: structural basis for homophilic adhesion via a novel dimerization motif. EMBO J 20, 4391–4398.

Kummer, D., Ebnet, K., 2018. Junctional Adhesion Molecules (JAMs): The JAM-Integrin Connection. Cells 7.

Kwan, K.M., Fujimoto, E., Grabher, C., Mangum, B.D., Hardy, M.E., Campbell, D.S., Parant, J.M., Yost, H.J., Kanki, J.P., Chien, C.B., 2007. The Tol2kit: a multisite gateway-based construction kit for Tol2 transposon transgenesis constructs. Dev Dyn 236, 3088–3099.

Lang, M.R., Patterson, L.B., Gordon, T.N., Johnson, S.L., Parichy, D.M., 2009. Basonuclin-2 requirements for zebrafish adult pigment pattern development and female fertility. PLoS Genet 5, e1000744.

Langer, H.F., Orlova, V.V., Xie, C., Kaul, S., Schneider, D., Lonsdorf, A.S., Fahrleitner, M., Choi, E.Y., Dutoit, V., Pellegrini, M., Grossklaus, S., Nawroth, P.P., Baretton, G., Santoso, S., Hwang, S.T., Arnold, B., Chavakis, T., 2011. A novel function of junctional adhesion molecule-C in mediating melanoma cell metastasis. Cancer Res 71, 4096–4105.

Larson, T.A., Gordon, T.N., Lau, H.E., Parichy, D.M., 2010. Defective adult oligodendrocyte and Schwann cell development, pigment pattern, and craniofacial morphology in puma mutant zebrafish having an alpha tubulin mutation. Dev Biol 346, 296–309.

Lewis, V.M., Saunders, L.M., Larson, T.A., Bain, E.J., Sturiale, S.L., Gur, D., Chowdhury, S., Flynn, J.D., Allen, M.C., Deheyn, D.D., Lee, J.C., Simon, J.A., Lippincott-Schwartz, J., Raible, D.W., Parichy, D.M., 2019. Fate plasticity and reprogramming in genetically distinct populations of Danio leucophores. Proc Natl Acad Sci U S A 116, 11806–11811.

Lopes, S.S., Yang, X., Muller, J., Carney, T.J., McAdow, A.R., Rauch, G.J., Jacoby, A.S., Hurst, L.D., Delfino-Machin, M., Haffter, P., Geisler, R., Johnson, S.L., Ward, A., Kelsh, R.N., 2008. Leukocyte tyrosine kinase functions in pigment cell development. PLoS Genet 4, e1000026.

Mahalwar, P., Singh, A.P., Fadeev, A., Nusslein-Volhard, C., Irion, U., 2016. Heterotypic interactions regulate cell shape and density during color pattern formation in zebrafish. Biology open 5, 1680–1690.

Mandicourt, G., Iden, S., Ebnet, K., Aurrand-Lions, M., Imhof, B.A., 2007. JAM-C regulates tight junctions and integrin-mediated cell adhesion and migration. J Biol Chem 282, 1830–1837.

McKeown, S.J., Wallace, A.S., Anderson, R.B., 2013. Expression and function of cell adhesion molecules during neural crest migration. Dev Biol 373, 244–257.

McMenamin, S.K., Bain, E.J., McCann, A.E., Patterson, L.B., Eom, D.S., Waller, Z.P., Hamill, J.C., Kuhlman, J.A., Eisen, J.S., Parichy, D.M., 2014. Thyroid hormone-dependent adult pigment cell lineage and pattern in zebrafish. Science 345, 1358–1361.

McMenamin, S.K., Chandless, M.N., Parichy, D.M., 2016. Working with zebrafish at postembryonic stages. Methods Cell Biol 134, 587–607.

Mills, M.G., Nuckels, R.J., Parichy, D.M., 2007. Deconstructing evolution of adult phenotypes: genetic analyses of kit reveal homology and evolutionary novelty during adult pigment pattern development of Danio fishes. Development 134, 1081–1090.

Milos, N.C., Wilson, H.C., Ma, Y.L., Mohanraj, T.M., Frunchak, Y.N., 1987. Studies on cellular adhesion of Xenopus laevis melanophores: modulation of cell-cell and cell-substratum adhesion in vitro by endogenous Xenopus galactoside-binding lectin. Pigment Cell Res 1, 188–196.

Morris, A.P., Tawil, A., Berkova, Z., Wible, L., Smith, C.W., Cunningham, S.A., 2006. Junctional Adhesion Molecules (JAMs) are differentially expressed in fibroblasts and co-localize with ZO-1 to adherens-like junctions. Cell Commun Adhes 13, 233–247.

Neiswender, J.V., Kortum, R.L., Bourque, C., Kasheta, M., Zon, L.I., Morrison, D.K., Ceol, C.J., 2017. KIT Suppresses BRAF(V600E)-Mutant Melanoma by Attenuating Oncogenic RAS/MAPK Signaling. Cancer Res 77, 5820–5830.

Nishimura, E.K., Yoshida, H., Kunisada, T., Nishikawa, S.I., 1999. Regulation of E- and P-cadherin expression correlated with melanocyte migration and diversification. Dev Biol 215, 155–166.

O’Reilly-Pol, T., Johnson, S.L., 2013. Kit signaling is involved in melanocyte stem cell fate decisions in zebrafish embryos. Development 140, 996–1002.

Parichy, D.M., 1996. Pigment patterns of larval salamanders (Ambystomatidae, Salamandridae): the role of the lateral line sensory system and the evolution of pattern-forming mechanisms. Dev Biol 175, 265–282.

Parichy, D.M., 2021. Evolution of pigment cells and patterns: recent insights from teleost fishes. Curr Opin Genet Dev 69, In press.

Parichy, D.M., Elizondo, M.R., Mills, M.G., Gordon, T.N., Engeszer, R.E., 2009. Normal table of postembryonic zebrafish development: staging by externally visible anatomy of the living fish. Developmental Dynamics 238, 2975–3015.

Parichy, D.M., Rawls, J.F., Pratt, S.J., Whitfield, T.T., Johnson, S.L., 1999. Zebrafish sparse corresponds to an orthologue of c-kit and is required for the morphogenesis of a subpopulation of melanocytes, but is not essential for hematopoiesis or primordial germ cell development. Development 126, 3425–3436.

Parichy, D.M., Reedy, M.V., Erickson, C.A., 2006. Chapter 5. Regulation of melanoblast migration and differentiation, in: Nordland, J.J., Boissy, R.E., Hearing, V.J., King, R.A., Oetting, W.S., Ortonne, J.P. (Eds.), The Pigmentary System: Physiology and Pathophysiology. 2nd Edition. Oxford University Press, New York, New York.

Parichy, D.M., Turner, J.M., 2003a. Temporal and cellular requirements for Fms signaling during zebrafish adult pigment pattern development. Development 130, 817–833.

Parichy, D.M., Turner, J.M., 2003b. Zebrafish puma mutant decouples pigment pattern and somatic metamorphosis. Developmental Biology 256, 242–257.

Patterson, L.B., Bain, E.J., Parichy, D.M., 2014. Pigment cell interactions and differential xanthophore recruitment underlying zebrafish stripe reiteration and Danio pattern evolution. Nat Commun 5, 5299.

Patterson, L.B., Parichy, D.M., 2013. Interactions with iridophores and the tissue environment required for patterning melanophores and xanthophores during zebrafish adult pigment stripe formation. PLoS Genet 9, e1003561.

Patterson, L.B., Parichy, D.M., 2019. Zebrafish Pigment Pattern Formation: Insights into the Development and Evolution of Adult Form. Annu Rev Genet 53, 505–530.

Patton, E.E., Mueller, K.L., Adams, D.J., Anandasabapathy, N., Aplin, A.E., Bertolotto, C., Bosenberg, M., Ceol, C.J., Chi, P., Herlyn, M., Holmen, S.L., Karreth, F.A., Kaufman, C.K., Khan, S., Kobold, S., Leucci, E., Levy, C., Lombard, D.B., Lund, A.W., Marie, K.L., Marine, J.C., Marais, R., McMahon, M., Robles-Espinoza, C.D., Ronai, Z.A., Samuels, Y., Soengas, M.S., Villanueva, J., Weeraratna, A.T., White, R.M., Yeh, I., Zhu, J., Zon, L.I., Hurlbert, M.S., Merlino, G., 2021. Melanoma models for the next generation of therapies. Cancer Cell.

Pinon, P., Wehrle-Haller, B., 2011. Integrins: versatile receptors controlling melanocyte adhesion, migration and proliferation. Pigment Cell Melanoma Res 24, 282–294.

Powell, G.T., Wright, G.J., 2011. Jamb and jamc are essential for vertebrate myocyte fusion. PLoS Biol 9, e1001216.

Powell, G.T., Wright, G.J., 2012. Genomic organisation, embryonic expression and biochemical interactions of the zebrafish junctional adhesion molecule family of receptors. PLoS ONE 7, e40810.

Provost, E., Rhee, J., Leach, S.D., 2007. Viral 2A peptides allow expression of multiple proteins from a single ORF in transgenic zebrafish embryos. Genesis 45, 625–629.

Quigley, I.K., Manuel, J.L., Roberts, R.A., Nuckels, R.J., Herrington, E.R., MacDonald, E.L., Parichy, D.M., 2005. Evolutionary diversification of pigment pattern in Danio fishes: differential fms dependence and stripe loss in D. albolineatus. `Development 132, 89–104.

Rao, C., Su, Z., Li, H., Ma, X., Zheng, X., Liu, Y., Lu, F., Qu, J., Hou, L., 2016. Microphthalmia-associated transcription factor regulates skin melanoblast migration by repressing the melanoma cell adhesion molecule. Exp Dermatol 25, 74–76.

Rathjen, F.G., 2020. The CAR group of Ig cell adhesion proteins-Regulators of gap junctions? Bioessays 42, e2000031.

Reymond, N., Garrido-Urbani, S., Borg, J.P., Dubreuil, P., Lopez, M., 2005. PICK-1: a scaffold protein that interacts with Nectins and JAMs at cell junctions. FEBS Lett 579, 2243–2249.

Roh-Johnson, M., Shah, A.N., Stonick, J.A., Poudel, K.R., Kargl, J., Yang, G.H., di Martino, J., Hernandez, R.E., Gast, C.E., Zarour, L.R., Antoku, S., Houghton, A.M., Bravo-Cordero, J.J., Wong, M.H., Condeelis, J., Moens, C.B., 2017. Macrophage-Dependent Cytoplasmic Transfer during Melanoma Invasion In Vivo. Dev Cell 43, 549–562 e546.

Santoso, S., Orlova, V.V., Song, K., Sachs, U.J., Andrei-Selmer, C.L., Chavakis, T., 2005. The homophilic binding of junctional adhesion molecule-C mediates tumor cell-endothelial cell interactions. J Biol Chem 280, 36326–36333.

Satohisa, S., Chiba, H., Osanai, M., Ohno, S., Kojima, T., Saito, T., Sawada, N., 2005. Behavior of tight-junction, adherens-junction and cell polarity proteins during HNF-4alpha-induced epithelial polarization. Exp Cell Res 310, 66–78.

Saunders, L.M., Aman, A.J., Mishra, A.K., Lewis, V.M., Toomey, M.B., Packer, J.S., Qiu, X., McFaline-Figueroa, J.L., Corbo, J.C., Trapnell, C., Parichy, D.M., 2019. Thyroid hormone regulates distinct paths to maturation in pigment cell lineages. eLife 8, e45181.

Schartl, M., Larue, L., Goda, M., Bosenberg, M.W., Hashimoto, H., Kelsh, R.N., 2016. What is a vertebrate pigment cell? Pigment Cell Melanoma Res 29, 8–14.

Scheiermann, C., Meda, P., Aurrand-Lions, M., Madani, R., Yiangou, Y., Coffey, P., Salt, T.E., Ducrest-Gay, D., Caille, D., Howell, O., Reynolds, R., Lobrinus, A., Adams, R.H., Yu, A.S., Anand, P., Imhof, B.A., Nourshargh, S., 2007. Expression and function of junctional adhesion molecule-C in myelinated peripheral nerves. Science 318, 1472–1475.

Schreiber, J., Langhorst, H., Juttner, R., Rathjen, F.G., 2014. The IgCAMs CAR, BT-IgSF, and CLMP: structure, function, and diseases. Advances in neurobiology 8, 21–45.

Shah, A.N., Davey, C.F., Whitebirch, A.C., Miller, A.C., Moens, C.B., 2015. Rapid reverse genetic screening using CRISPR in zebrafish. Nat Methods.

Sharan, S.K., Thomason, L.C., Kuznetsov, S.G., Court, D.L., 2009. Recombineering: a homologous recombination-based method of genetic engineering. Nat Protoc 4, 206–223.

Singh, A.P., Dinwiddie, A., Mahalwar, P., Schach, U., Linker, C., Irion, U., Nusslein-Volhard, C., 2016. Pigment Cell Progenitors in Zebrafish Remain Multipotent through Metamorphosis. Dev Cell 38, 316–330.

Solnica-Krezel, L., Schier, A.F., Driever, W., 1994. Efficient recovery of ENU-induced mutations from the zebrafish germline. Genetics 136, 1401–1420.

Spiewak, J.E., Bain, E.J., Liu, J., Kou, K., Sturiale, S.L., Patterson, L.B., Diba, P., Eisen, J.S., Braasch, I., Ganz, J., Parichy, D.M., 2018. Evolution of Endothelin signaling and diversification of adult pigment pattern in Danio fishes. PLoS Genet 14, e1007538.

Steinbacher, T., Kummer, D., Ebnet, K., 2018. Junctional adhesion molecule-A: functional diversity through molecular promiscuity. Cell Mol Life Sci 75, 1393–1409.

Suster, M.L., Abe, G., Schouw, A., Kawakami, K., 2011. Transposon-mediated BAC transgenesis in zebrafish. Nat Protoc 6, 1998–2021.

Suster, M.L., Kikuta, H., Urasaki, A., Asakawa, K., Kawakami, K., 2009. Transgenesis in zebrafish with the tol2 transposon system. Methods Mol Biol 561, 41–63.

Takahashi, G., Kondo, S., 2008. Melanophores in the stripes of adult zebrafish do not have the nature to gather, but disperse when they have the space to move. Pigment Cell Melanoma Res 21, 677–686.

Twitty, V.C., 1945. The developmental analysis of specific pigment patterns. Journal of Experimental Zoology 100, 141–178.

Ustun, Y., Reibetanz, M., Brachvogel, B., Nischt, R., Eckes, B., Zigrino, P., Krieg, T., 2019. Dual role of laminin511 in regulating melanocyte migration and differentiation. Matrix Biol 80, 59–71.

Usui, Y., Aramaki, T., Kondo, S., Watanabe, M., 2019. The minimal gap-junction network among melanophores and xanthophores required for stripe pattern formation in zebrafish. Development 146.

Watanabe, M., Iwashita, M., Ishii, M., Kurachi, Y., Kawakami, A., Kondo, S., Okada, N., 2006. Spot pattern of *leopard Danio* is caused by mutation in the zebrafish connexin41.8 gene. EMBO Rep 7, 893–897.

Watanabe, M., Kondo, S., 2015. Is pigment patterning in fish skin determined by the Turing mechanism? Trends Genet 31, 88–96.

Watanabe, M., Sawada, R., Aramaki, T., Skerrett, I.M., Kondo, S., 2016. The Physiological Characterization of Connexin41.8 and Connexin39.4, Which Are Involved in the Striped Pattern Formation of Zebrafish. J Biol Chem 291, 1053–1063.

Williams, J.S., Hsu, J.Y., Rossi, C.C., Artinger, K.B., 2018. Requirement of zebrafish pcdh10a and pcdh10b in melanocyte precursor migration. Dev Biol 444 Suppl 1, S274–S286.

